# Bacterial Strategies for Damage Management

**DOI:** 10.1101/2022.10.27.514133

**Authors:** Kunaal Joshi, Rhea Gandhi, David C. Krakauer, Srividya Iyer-Biswas, Christopher P. Kempes

## Abstract

Organisms are able to partition resources adaptively between growth and repair. The precise nature of optimal partitioning and how this emerges from cellular dynamics including insurmountable trade-offs remains an open question. We construct a mathematical framework to estimate optimal partitioning and the corresponding maximal growth rate constrained by empirical scaling laws. We model a biosynthesis tradeoff governing the partitioning of the ribosome economy between replicating functional proteins and replicating the ribosome pool, and also an energy tradeoff arising from the finite energy budget of the cell. Through exact analytic calculations we predict limits on the range of values partitioning ratios take while sustaining growth. We calculate how partitioning and cellular composition scale with protein and ribosome degradation rates and organism size. These results reveal different classes of optimizing strategies corresponding to phenotypically distinct bacterial lifestyles. We summarize these findings in a quadrant-based taxonomy including: a “greedy” strategy maximally prioritizing growth, a “prudent” strategy maximally prioritizing the management of damaged pools, and “strategically limited” intermediates.

## I. INTRODUCTION

A fundamental question for all of life is how to manage damage accumulation while balancing resources between repair and growth, i.e., how to partition metabolic resources between biosynthesis and maintenance. Recently a variety of efforts have shown the importance of maintenance considerations for predicting growth across diverse species [1–6]. In such efforts maintenance has been shown to necessitate a constant energetic cost per unit volume, independent of the size of the organism. The cost for maintenance has also been shown to be a complicated sum of the replacement costs associated with multiple different pathways and macromolecules. However, it is unclear how these costs emerge from an evolutionary optimization process when accounting for trade-offs amongst physiological functions. This is material since changes in maintenance costs significantly influence growth rates at a level that can be detected by selection [4, 6–8]. Thus, for any replacement process, a key question is what level of investment is optimal for fitness.

We provide a mathematical framework for calculating the optimization of growth rate when constrained by the energetic costs of eliminating damage and the growth-inhibiting consequence of allowing damage to accumulate. In this paper we focus on the coupled system of ribosomes and functional proteins. Our framework couples two previous theories for cell growth which both make predictions across species based on cell size [4, 5].

We find that there is a well-defined global minimum division time for cells in the space of possible energy and ribosome allocations. Significantly, for *E. coli*, this minimum corresponds to the maximum growth rates observed for this organism. Furthermore, the isoclines of fitness around the global minimum indicate that evolutionary trajectories of neutral fitness take on complicated inter-dependencies.

We also considered how the global minimum changes with cell size across species, since our framework is based on coupling two models that are fundamentally scale dependent. We find that the fraction of biosynthesis dedicated to producing ribosomes increases with cell size, and that the energy allocated to growth also increases. Taken together this leads to joint optimum where division time decreases with cell size consistent with previous observations [4, 9]. We also find that this shift in the optimum allocation of resources, and the corresponding division time, qualitatively trends in the opposite direction with increasing degradation rate as when the cell size increases. Thus, the effect of degradation rate on growth dynamics decreases as cell size increases.

## II. THE OPTIMIZATION FRAMEWORK

Cellular damage processes range from decay of macromolecules to “writing” errors during transcription, translation, and DNA copying. In each case there are many possibilities for how damage could affect cellular physiology. For example, damaged products could: (i) interact with other cellular components in a way that damages those components; (ii) decrease overall cellular rates such as biosynthetic production rates; (iii) increase the “error” rate of the production processes producing functional or damaged products; or (iv) simply displace functional components.

This is a rich space of possibilities. Here we focus on possibility (iv) since it is fundamental and an inescapable consequence of damage. Specifically, we consider a scenario in which damaged products take up physical space and require energy to be removed in order to free up space for functional products. In this way we deal with both the material and energy economies of the cell. A schematic of our optimization framework is provided in Figure 1. It should be noted that possibilities (i-iii) could be added to more complex models, but we would argue that (iv) is always a consideration.

**FIG. 1:**
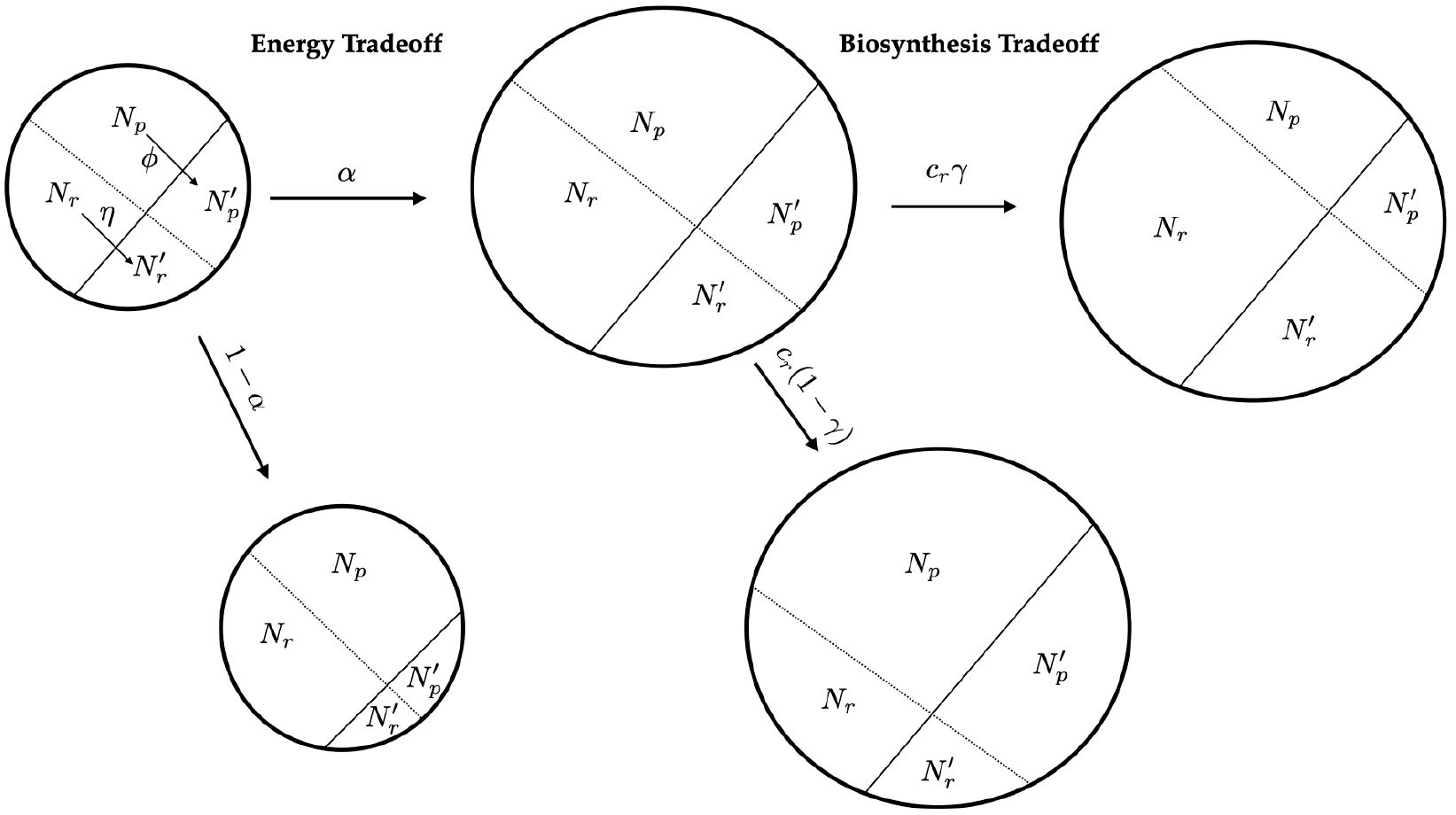
Schematic representation of framework for growth/energy and biosynthesis trade-offs. Cell volume is accounted for by functional ribosomes (*N_r_*) and proteins (*N_p_*), and damaged ribosomes 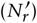 and proteins 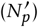. Functional ribosomes and proteins degrade into damaged ribosomes and proteins at constant rates *η* and *ϕ*. Energy production of the cell is determined by the number of functional proteins *N_p_* and split between between growth (production of new ribosomes and proteins) and replacement of damaged ribosomes and proteins in the ratio *α* : 1 – *α*. The energy spent on replacement is used to clear out damaged ribosomes and proteins, freeing up cell volume that can be be occupied by functional ribosomes and proteins speeding up cell growth. New proteins and ribosomes are produced by the existing ribosomes at the rate *c_r_* (polymerization rate) that is determined by the energy budget. Out of the total biosynthesic capacity of the existing ribosomes, a fraction *γ* is allocated to ribosome production, and the remaining 1 – *γ* to protein production. Cell division occurs when the cell volume exactly doubles since the last division. The fastest growing population is comprised of cells with optimum partitioning ratios *γ* and *α* such that division time is minimized at steady state.

### The ribosome economy

Previous work has shown that the number of ribosomes required by a functioning cell can be predicted by considering the dynamics of replicating the ribosome and protein pools given a division time and a required number of expressed proteins [5, 10–12]. In this approach, the total biosynthetic capacity (determined by the polymerization rate *c_r_* of the ribosome) is split into ribosome production and protein production according to *γ* and 1 – *γ* respectively, such that both protein and ribosome pools double in the duration of a cell cycle under ideal circumstances. This assumption ensures that physiology is consistent across generations. The functional ribosome (*N_r_*) and protein numbers (*N_p_*) in the cell thus evolve as follows:

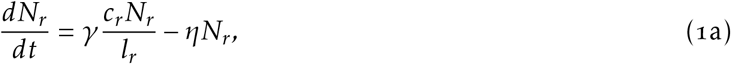

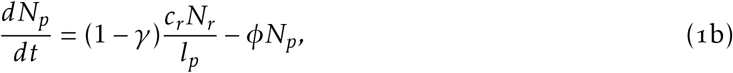

wherein the last terms in both equations are the degradation (error rates) rates for ribosomes and proteins, and *l_r_* and *l_p_* are the average ribosome and protein codon lengths. Since we are interested in the tradeoffs associated with investing in replacing damaged products, we explicitly track the transfer of ribosomes and proteins into damaged pools with populations 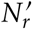 and 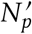. The production rates of these damaged populations are equal to the ribosome or protein degradation rates, minus the rate at which these degraded substances are removed from the system (the damaged ribosomes and proteins removed are labeled 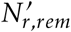 and 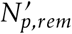):

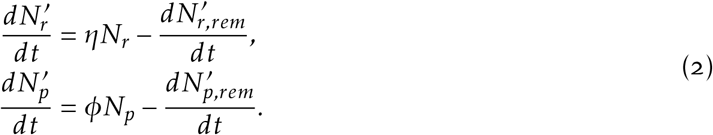

Solving Eqs. 1 and 2, we obtain the time evolution of ribosome and protein numbers:

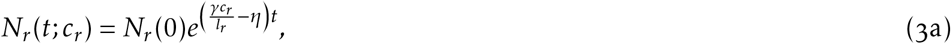

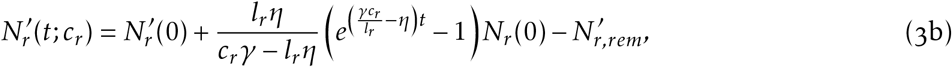

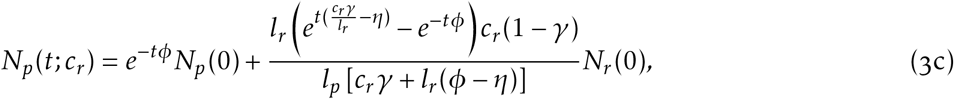

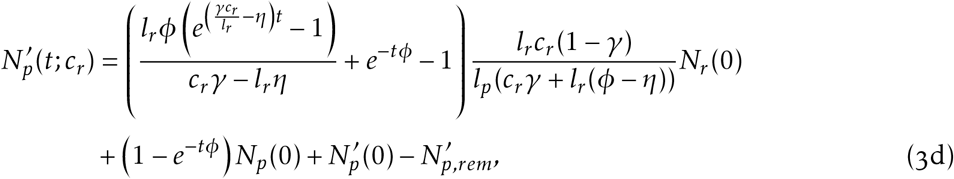

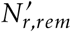 and 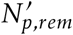 are the number of damaged ribosomes and proteins cleared by the cell, which depend on the energy allocated to the removal of damaged pools and the cell’s strategy for clearing damaged ribosomes vs. damaged proteins as described in detail below.

### The energy budget

The foregoing equations give growth dynamics following only biosynthetic constraints. However, growth is also fundamentally limited by energetic constraints and it is important to couple biosynthesis with metabolism. We assume that the instantaneous energy production follows a scaling relationship with cell size that ultimately depends on the protein abundance as follows:

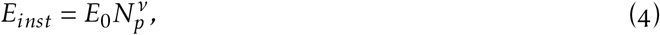

where *v* and *E*_0_ are calculated from interspecific data (see Supplementary Section VI B)[4, 9]. The total energy produced during a division cycle, over a division period *τ*, is given by

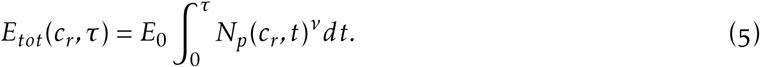

Out of this total energy, a fraction *α* is allocated to growth and 1 – *α* to removal of damaged pools. Production of new proteins and ribosomes requires energy. The total energy required (*E_req_*) during a division cycle is given by integrating the production rates of ribosomes and proteins over the division cycle and multiplying by the energetic costs of producing a new unit,

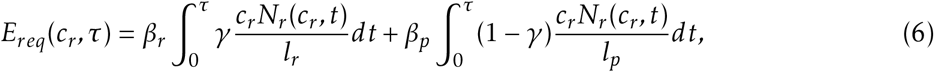

where *β_r_* is the energetic cost of producing one ribosome, and *β_p_* is the energetic cost of producing one protein. From Eq. 1 and 2,

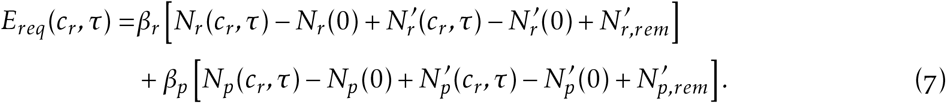

Thus growth is directly limited by the energy allocated to growth. We incorporate this effect by choosing the highest possible polymerization rate, *c_r_*, such that the energy allocated to growth is sufficient to cover the energetic costs of ribosome and protein production over an entire cell cycle. Thus, for a given division time *τ*, *c_r_* is chosen to be the maximum possible value that satisfies

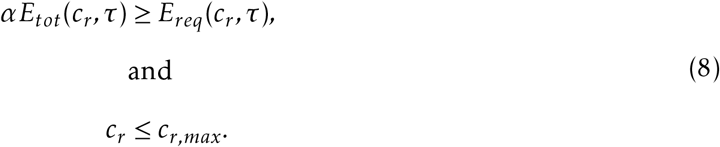

Here, we impose a physiological limit to the maximum achievable rate of polymerization, *c_r_,_max_*, beyond which *c_r_* cannot rise even if there is excess energy available.

As damaged ribosomes and proteins accumulate in the cell, there is less volume available for functional ribosomes and proteins, which negatively impacts cell growth. Thus, cells spend the remaining fraction 1 – *α* of the total energy for clearing out damaged pools and freeing up volume for functional pools. The energy required per unit volume to clear damaged ribosomes is less than that required to clear damaged proteins, thus we assume a cellular strategy of spending the energy allocated initially only for clearing damaged ribosomes. If there is excess energy left after the damaged ribosomes are completely cleared, that energy is spent on clearing damaged proteins.

The portion of the total cell volume allocated to ribosomes and proteins is given by

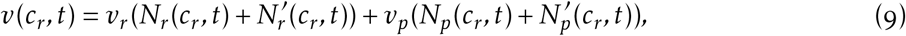

where *v_r_* is the average volume occupied by one ribosome, and *v_p_* is the average volume occupied by one protein. We assume that the total volume of the cell is proportional to the volume *v* given by the sum of the volumes occupied by the ribosome and protein pools and the damaged ribosome and protein pools (see supplementary section VI B2).

We take the division time to be the time taken to exactly double the cell volume (i.e., we assume a deterministic model of a cellular division where the total cell volume is partitioned equally among daughter cells upon division). Once cells reach steady state, each of the pools exactly doubles in volume in the duration of one division cycle (i.e., *N_r_*(*c_r_,τ*) = 2*N_r_*(*c_r_*, 0) etc.). Adding this condition to Eq. 3a, we get the following relation between division time, *τ*, and *c_r_*,

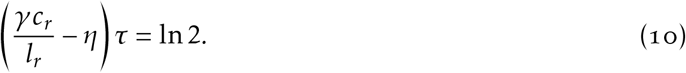

Using this to eliminate *τ* from Eqs. 3a–3d, the steady state initial values of *N_r_*, 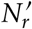 and 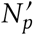 can be now expressed in terms of initial value of *N_p_*, the polymerization rate *c_r_*, and other fixed quantities (see the supplementary section VI E for details). Incorporating these expressions in Eq. 8, we arrive at

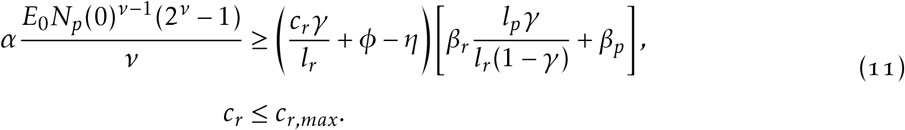

Thus, the maximum value of *c_r_* can be expressed in terms of initial value of *N_p_.*

Note that although this constraint ensures that the total energy produced throughout the division cycle would be sufficient to support growth, it doesn’t ensure that the instantaneous energy produced at any instant in the division cycle would be greater than or equal to the instantaneous energy required for growth at that instant. To overcome this, our model requires the implicit assumption that the cell has an energy bank, such as ATP pools, that it manages over a lifecycle.

It is important to note that *N_p_* (0) can itself be fixed using the prescribed initial volume, *v*_0_, of the cell:

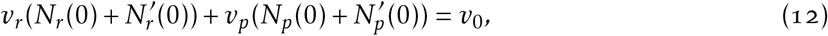

where all terms on LHS can be written in terms of *N_p_*(0). Substituting the value of *N_p_*(0), obtained by numerically solving this implicit equation, into Eqs. 11 and 10, we obtain the value of steady state division time for a given *γ* and *α*. We find the optimum values of *γ* and *α* which minimize the division time by varying *γ* and *α* over the entire range of possible values and repeating the optimization procedure just described.

In steady state, both functional ribosome pool and functional protein pool grow exponentially at the same rate (see Eq. 3a and supplementary Eq. 34) given by *γc_r_/l_r_* – *η*. The damaged ribosome and protein pools also exactly double in the same duration, but their specific growth function is dependent on how the damaged pools are cleared. In our model, the total damaged ribosomes and proteins removed are given by the total energy allocated to the removal of damaged pools, but our model is agnostic to the specific schedule for how this energy is spent as a function of time over the division cycle. Thus, through an appropriate choice of removal as a function of time, damaged pools can be made to also grow exponentially at the same rate (ensuring total cell volume grows exponentially with a constant rate given by *γc_r_/l_r_* – *η*, or, equivalently, (ln2)/*τ*), although this is not a strict requirement of our model.

In summary, we are optimizing two trade-offs through two control parameters. The first involves splitting the total available biosynthetic capacity between ribosome and protein production (optimizing *γ*). Increasing ribosome production increases the biosynthetic capacity, while increasing protein production increases the energy available for growth and the removal of damaged pools, thus a balance between the two is required to optimize overall growth. The second trade-off involves splitting the available energy between growth of protein and ribosome pools and the removal of damaged protein and ribosome pools (optimizing *α*). Since total cell volume is limited, accumulation of damaged protein and ribosome pools takes away viable space from the functional protein and ribosome pools, thereby leading to a decrease in growth rate and cell fitness. Thus, it is necessary to balance energy spent on growth with energy spent on the removal of damaged pools.

## III. RESULTS

### A. Optimal Growth

We first quantified the trade-off in biosynthesis and energy partitioning for *E. coli* cells (Fig. 2). Ribosomes are the source of a cell’s biosynthesis capacity, while expressed proteins are responsible for generating energy. Optimizing production of new ribosomes and proteins requires the strategic allocation of both energy and biosynthesis capacity in an appropriately balanced amount. For low *γ* values, cells wouldn’t have sufficient biosynthetic ability to support fast growth, while at high *γ* values, the cell’s growth would be limited by energy. This trade-off can be seen in Fig. 2, where growth is not viable for *γ* < 0.0045 due to insufficient biosynthetic capacity, and growth slows (division time increases) as the system approaches this limit (Fig. 2A i). Furthermore, as *γ* values increase, the minimum value of *α* required for viable growth increases, indicating the minimum energy that the cell needs to allocate to growth rather than the removal of damaged pools increases due to energy becoming limiting at higher *γ* values (Fig. 2A ii).

**FIG. 2:**
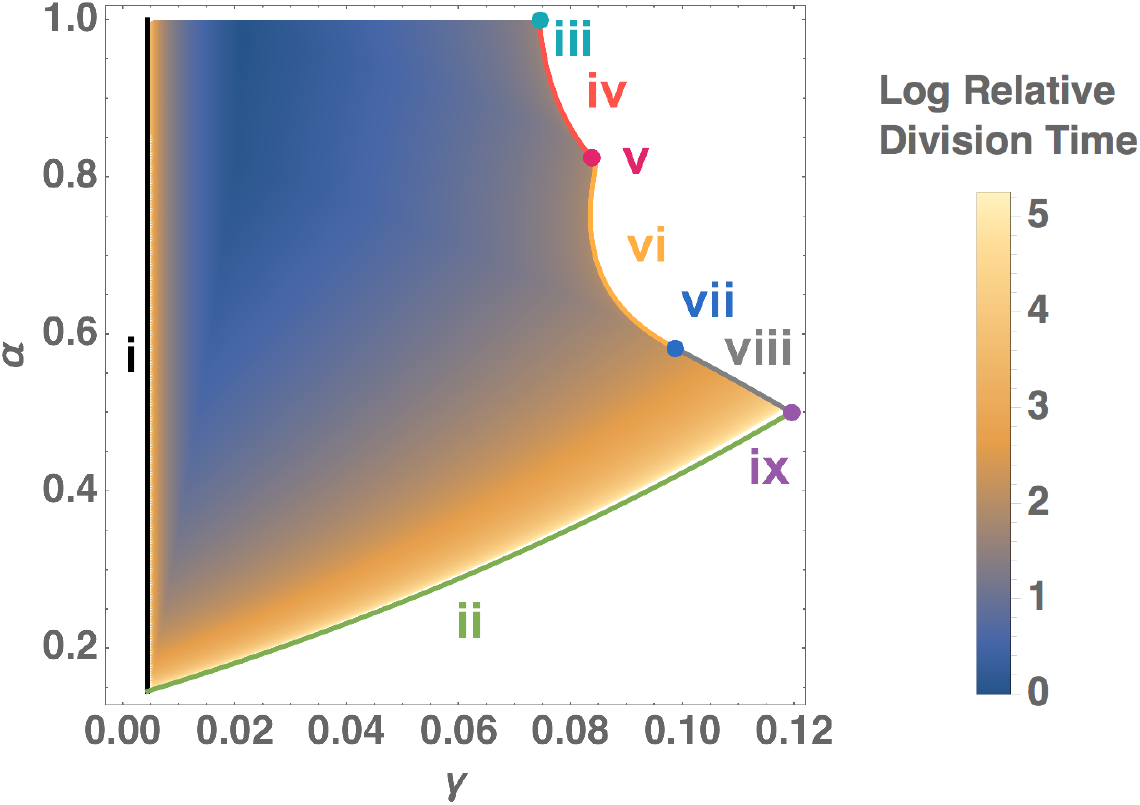
Trade-offs in energy and biosynthesis partitioning. *γ* is the fraction of total biosynthesis allocated to ribosome production (vs protein production), and *α* is the fraction of total energy allocated to growth (vs the removal of damaged pools). The heatmap shows the variation of steady state division time with *γ* and *α*, over the full range of values. The theoretical bounds on *γ* and *α* denoted by i-ix are derived in supplementary section VI F. There exist multiple regions of local minima in this heatmap, but we focus on the global minimum. The global minimum is 2993.44 (s) at *γ* = 0.0213 and *α* = 0.9766, when measured with a least count of 0.0001 for both *γ* and *α*. Division time is plotted relative to the global minimum division time.

In conjunction with optimizing biosynthesis partitioning, the cell also needs to optimize its energy expenditure between growth and the removal of damaged pools. For cells that employ a “*prudent*” strategy of prioritizing damage control (low *α* limit), cell growth is limited simply by the lack of energy allocated to growth. At the other extreme, cells employing a “*greedy*” strategy of allocating the maximum resources to growth (high *α* limit) face the issue of a significant fraction of their volume becoming unusable due to accumulating damaged pools. This limits the energy production and biosynthetic capacity of the cell, which in turn limits cell growth. On increasing *γ*, cells are effectively increasing their biosynthetic capacity in exchange for reduced energy production, which increases the minimum *α* required for viable growth to ensure cells have at least the bare minimum energy for growing. This places a limit on how *“prudent”* the cells are allowed to be while still sustaining viable growth. Importantly, beyond a certain value of *γ* (Fig. 2A iii), we found that it becomes essential for cells to also allocate a minimum amount of energy towards the removal of damaged pools to ensure that cells don’t accumulate damage each generation, eventually leading to cell death without reaching steady state, essentially placing a physiological limit on the “*greedy*” strategy. This upper limit on *α* decreases as *γ* increases (Fig. 2A iv), indicating the need to allocate more energy to clearing damaged ribosomes, and eventually reaching a point where all damaged ribosomes are cleared (Fig. 2A v). On increasing *γ* beyond this point, additional energy needs to be allocated towards the removal of damaged proteins (Fig. 2A vi). Beyond a certain *γ* value (Fig. 2A vii), both damaged ribosomes and proteins need to be fully cleared to sustain growth. The minimum energy fraction needed to achieve this increases (i.e., upper bound on *α* decreases) as *γ* increases (Fig. 2A viii), until this upper bound intersects with the lower bound on *α* (Fig. 2A ix). Growth is not viable for any *γ* value above this limit. The analytic solutions for bounds i-ix are derived in supplementary section VI F.

Within the *γ* – *α* phase space bounded by the above limits (Fig. 2A), cells face two trade-offs: (i) allocation of biosynthesis to producing ribosomes (to increase biosynthesis availability) vs proteins (to increase energy availability), and (ii) allocation of energy to growth vs removal of damaged pools. We find that the absolute global minimum for division time, for cells of size comparable to *E. coli*, and degradation rates comparable to *E. coli’s* base rates, is 2993.64 (s) at optimum allocation ratios *γ* = 0.0213 and *α* = 0.9766 (Fig. 2B). This lies in the intermediate *“strategically limited*” region between the “*prudent*” and “*greedy*” strategies, but closer to the “*greedy*” strategy for this particular case. At these parameter values, the damaged ribosome pool is completely cleared, while damaged proteins make up 27% of the total protein pool in steady state. For comparison, ribosomes occupy 21% of the total volume occupied by ribosomes and proteins.

### B. Effects Cell Size and Degradation Dynamics

As mentioned earlier, both cell size and degradation rates significantly affect the growth rate of cells and the optimal ratios of macromolecules [4, 5, 9]. Within the context of our model we can address the consequences of both.

The optimum partitioning at different degradation rates, while keeping the average cell size constant, is shown in Fig. 3A-C. We find that both the optimal *γ* and *α* decrease with increasing degradation rates while the achievable growth rate decreases. As the degradation rate increases, cells accumulate larger damaged protein and ribosome pools, and the importance of spending energy on clearing these pools increases accordingly. Consequently, there is a shift in the cell’s optimal strategy from *“greedy”* to *“prudent”*, marked by a decrease in the optimum *α*. This also manifests as a decrease in optimum *γ*, indicating cells are allocating more biosynthetic capacity to producing proteins which are responsible for generating energy in the cells.

**FIG. 3:**
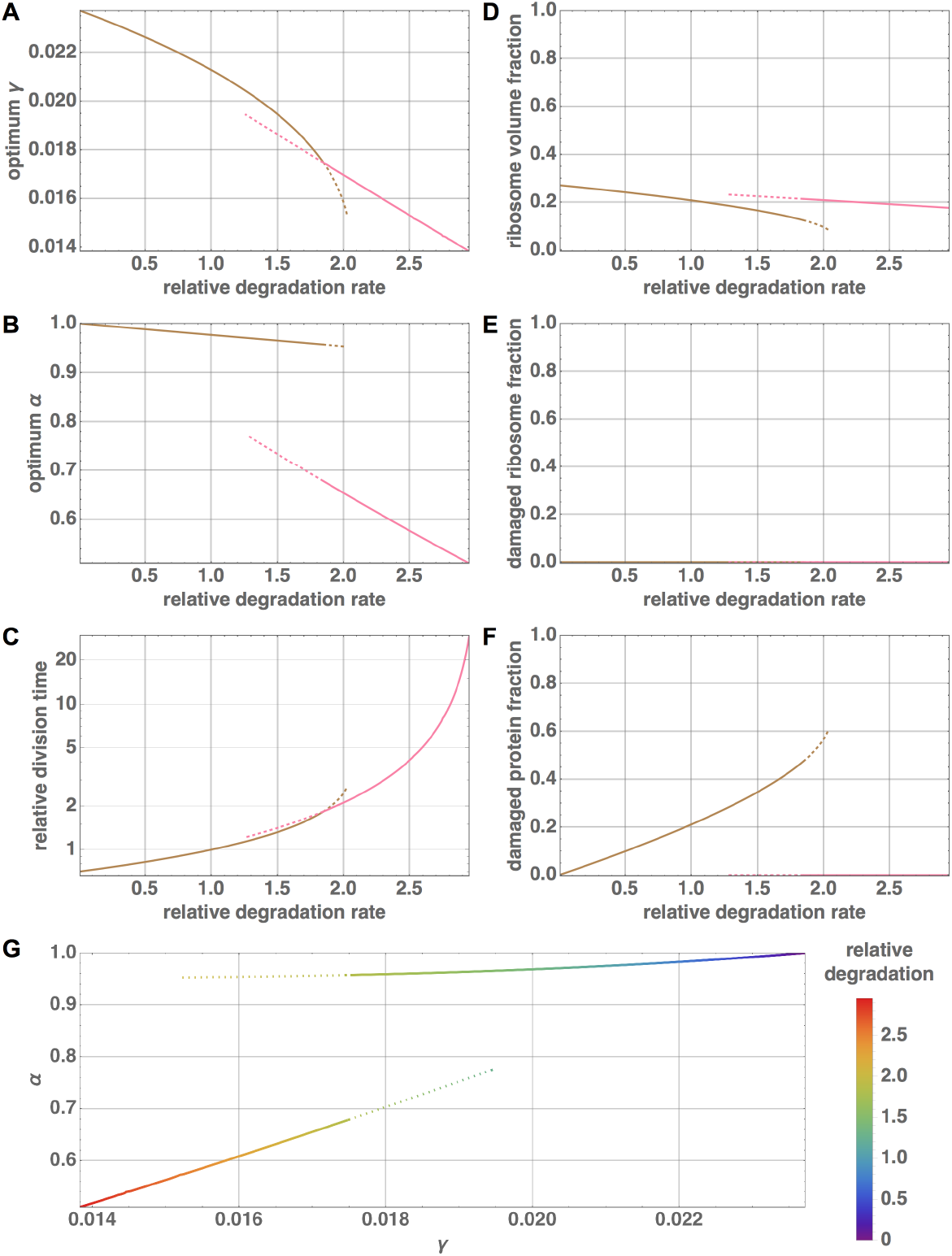
Scaling of optimum partitioning with degradation rate. A-C: Variation of optimum *γ* (A), *α* (B), and division time (C) with degradation rate (the degradation rates of ribosomes and proteins are taken to be equal, and are normalized by the base degradation rate of *E. coli*). The size is kept constant at the size of *E. coli*. For a given degradation rate, the values of *γ* and *α* at which the division time is minimum, and this minimum division time is plotted. Division time is measured relative to the minimum division time at base degradation rate of *E. coli*. Solid lines represent the global minima, and dotted lines represent local minima. The brown minima lie in the regime in which cells clear out all damaged ribosomes, but not all damaged proteins, while the pink minima lie in the regime in which cells clear out all damaged ribosomes and proteins. D-F: The division of cell volume between functional and damaged ribosomes and proteins at the optimum partitioning ratios is plotted as a function of degradation rate. D: Ribosome volume fraction is the fraction of total ribosome and protein volume occupied by functional and damaged ribosomes. E: Damaged ribosome fraction is the fraction of total ribosomes composed of damaged ribosomes. F: Damaged protein fraction is the fraction of total proteins composed of damaged proteins. G: The trajectories traced by the minima in *γ* – *α* space as degradation rate changes.

The optimum partitioning at different cell sizes while keeping the degradation rate constant is shown in Fig. 4A-C. For a given cell size we calculate the initial volume occupied by ribosome and protein pools (*v*_0_) and energy production constant (*E*_0_) using empirical scaling data (see supplementary section VI B 2 for more details). As size increases, optimum *γ* significantly increases, indicating cells allocate significantly more resources (biosynthesis and energy) towards ribosomes compared to proteins. This matches previously known trends of preference towards ribosomes at larger sizes [5]. Optimum *α* also increases with size, indicating a shift from *“prudent”* to “*greedy*”, and ultimately becomes 1 for sizes above a certain threshold.

**FIG. 4:**
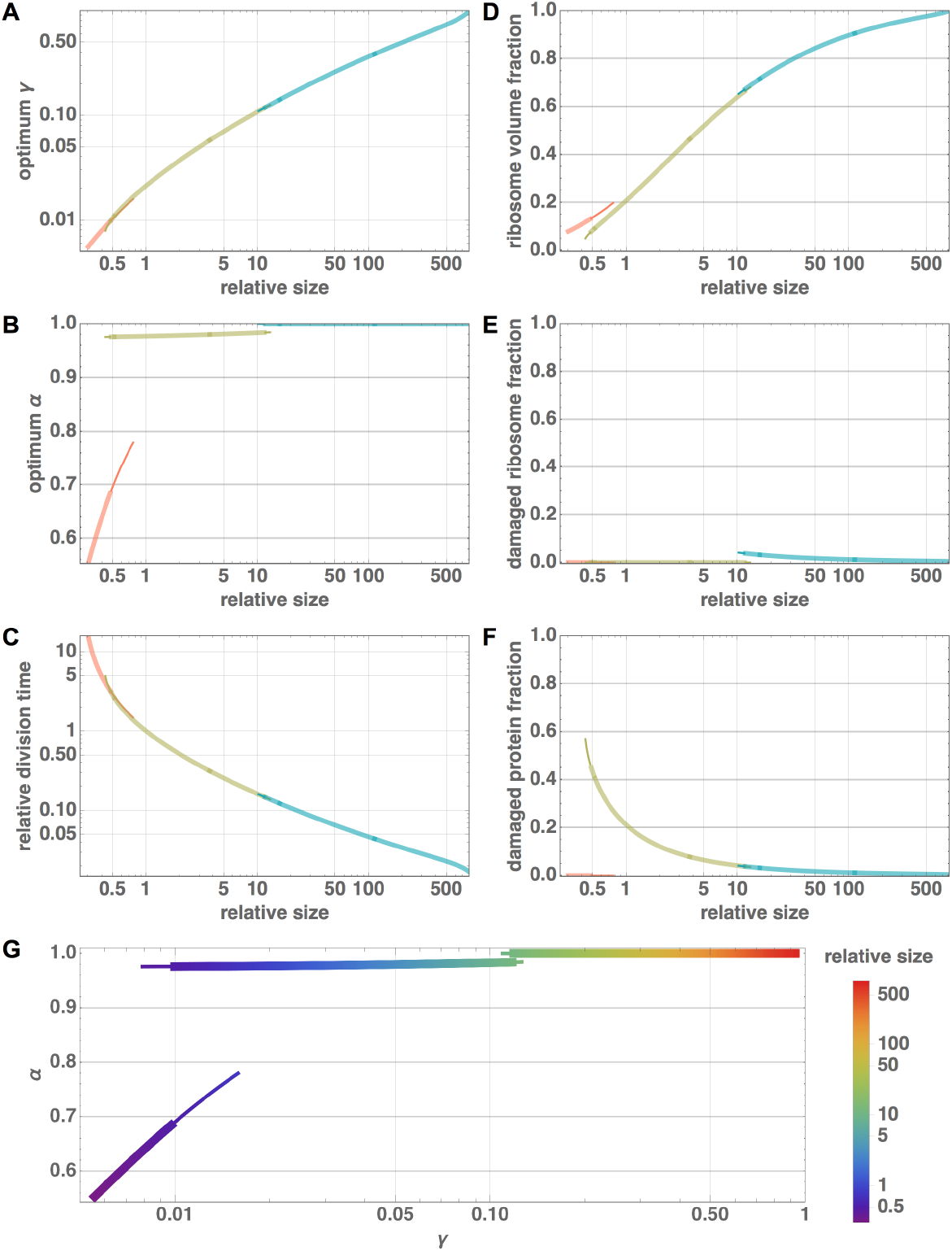
Scaling of optimum partitioning with cell size. A-C: Variation of optimum *γ* (A), *α* (B), and division time (C) with cell size (normalized by the size of *E. coli).* Degradation rate is kept constant at base degradation rate of *E. coli.* For cells of a given average size, the values of *γ* and *α* at which the division time is minimum, and this minimum division time are plotted. Division time is measured relative to the minimum division time at *E. coli* cell size. Thick lines represent the global minima, and thin lines represent local minima. The three minima lie in distinct regimes: (i) red: all damaged ribosomes and proteins are cleared, (ii) green: damaged ribosomes are fully cleared but not damaged proteins, and (iii) blue: neither damaged ribosomes nor damaged proteins are fully cleared. D-F: The division of cell volume between functional and damaged ribosomes and proteins at the optimum partitioning ratios is plotted as a function of cell size. D: Ribosome volume fraction is the fraction of total ribosome and protein volume occupied by functional and damaged ribosomes. E: Damaged ribosome fraction is the fraction of total ribosomes composed of damaged ribosomes. F: Damaged protein fraction is the fraction of total proteins composed of damaged proteins. G: The trajectories traced by the minima in *γ* – *α* space as cell size changes.

On varying size or degradation rate, we observe different division time local minima in the *γ* – *α* space representing qualitatively different choices, ranging from the most *“prudent”* to the most “*greedy*”: whether to allocate enough energy to completely clear out both damaged pools, just damaged ribosomes, or neither. For a given degradation rate and cell size, we observe that the local minimum with the largest *γ* generally turns out to be the global minimum, consistent with Eq. 10 which shows an inverse relationship between *γ* and division time, though this observation doesn’t follow directly from that equation (because the *c_r_* in that equation is also dependent on *γ* in a complex manner). Thus, the transitions in global minima coincide with the intersection of gamma values with the different local minima. The various local minima eventually either enter the basin of attraction of the corresponding global minimum and vanish, or cross the bounds for viable growth and vanish.

At low degradation rates, the most optimum partitioning corresponds to the local minimum where all damaged ribosomes are cleared, but not all damaged proteins. As degradation rate increases, the most optimum partitioning transitions to the regime in which all damaged ribosomes and proteins are cleared (the most *“prudent”* strategy), marked by a sharp decrease in *α*, i.e., more energy allocated to clearing damaged pools (Fig. 3). For size scaling, the reverse trend is observed compared to degradation scaling. As size increases, more volume availability for functional protein and ribosome pools seems to effectively reduce the effect of a constant degradation rate on overall growth. For small sizes, the optimum partitioning corresponds to the local minimum with both damaged pools completely cleared (which is also observed at high degradation rates, corresponding to the most *“prudent”* strategy). On increasing size, the optimum transitions to the local minimum where damaged ribosomes are completely cleared, and subsequently transitions to the local minimum where neither pools are completely cleared (Fig. 4). Eventually, the optimum further transitions to the regime in which all available energy is solely allocated to growth (*α* = 1, the most *“greedy”* strategy). Thus, the effect of increasing size on optimum partitioning and growth rate is qualitatively similar to decreasing the degradation rate. In Fig. 3, for scaling with degradation with cell size set to the size of an *E. coli*, we do not see any local minima in the regime in which neither damaged pools are fully cleared because the cell size is not large enough for this regime to have a local minimum for any degradation rate.

A major prediction of previous work on bacterial scaling is the dependence of growth rate on cell size [4, 9]. Since these previous efforts are based on the same energetic constraints used here, but without the additional feedback with the functional proteins nor the constraints of biosynthesis, it is worth exploring how growth rate depends on cell size in our framework here. Fig. 4C shows the dependence of optimal division time on cell size. For large cell sizes, we find that division time scales with cell volume according to an exponent of −0.63, which compares well with the best fit to data of −0.64 [4, 9]. Since our model is based on the connection between growth, protein abundances, and size it will also be interesting to connect these results to the known shifts in cell size with protein production [13]. Similarly, a variety of recent work has investigated the detailed statistics of cellular growth and division in order to uncover the underlying mechanisms of cellular homeostasis [14–19] and it will also be interesting to connect those detailed mechanisms with these size-based results.

## IV. CONCLUSION

Maintenance and replacement investment is a key determinant of the growth of any organism [1, 4]. Bacterial cells make complicated trade-offs associated with investing in the removal of damaged pools of biomolecules so as to optimize growth. Here we focused on a key aspect of cellular physiology, the replication of functional proteins and ribosomes, to show how maximal growth rate emerges as the natural optimization of energetic and biosynthetic partitioning. We showed that for a fixed cell size a nontrivial global optimum exists and that this optimum is consistent with maximum growth rates in *E. coli*. We also show that across cells of different size the allocations to growth — both the biosynthesis dedicated to ribosome production and energetic allocation to growth — increase with increasing cell size. This leads to a scaling of division time with cell size that impressively matches previous observations and theory [4, 9].

Using these results we are able to classify microbial growth and repair strategies into four functional varieties in a larger trade-off space (Fig. 5). Two quadrants experience opposing pressures on their control parameters. Large cells that experience significant damage and small cells that experience minimal damage. Both of these life histories find themselves caught between maximizing growth and maximizing maintenance and unable to specialize on either. And two quadrants experience aligned constraints allowing them to either maximize growth - large cells with low damage - or maximize maintenance - small cells with high damage. Excitingly, these position us to ask how these predictions are born out in the life history of bacterial lineages.

**FIG. 5:**
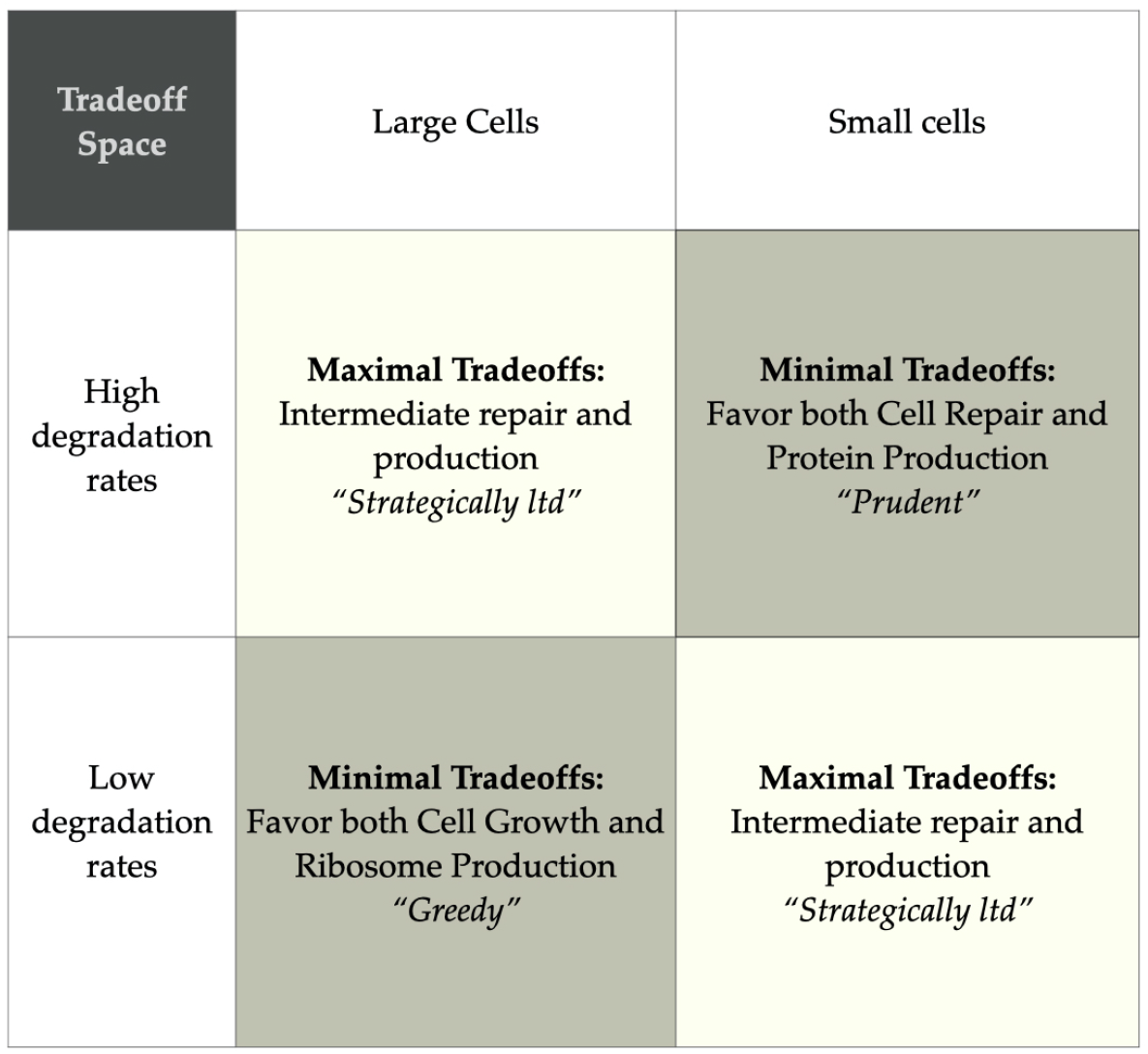
Functional Strategies in Bacterial Trade-off Space. The above table qualitatively summarizes the optimal strategies for fastest growth in different regimes of degradation rates and cell sizes. Increasing degradation rate tips the scales towards the more *“prudent”* strategies, while increasing cell size conversely favors *“greedy”* strategies. For large cells with adequately high degradation rates, or small cells with correspondingly low degradation rates, an intermediate strategy is preferred, and we see the maximal trade-offs for repair and production partitioning.

Looking ahead this framework may be a powerful tool for assessing physiological tradeoffs in a variety of environments and understanding evolutionary trajectories. For example, in contexts where the degradation rate of proteins is shifted due to environmental stresses or antibiotics from competing organisms our framework could be used to assess the new optimal growth rate and predict evolutionary or physiological responses.

Our work also highlights the importance of including damage and the removal of damaged pools in considerations of bacterial physiology. Other recent work has highlighted the importance of including degradation in growth models in order to accurately predict the ribosomal mass fraction at slow growth [12] where there is curvature away from the linear “growth law” [10]. Most of these efforts, along with the ribosome model that we employed here [5], focus on the physiological consequences associated with a particular growth rate. Here we have expanded on this work to optimize growth under various physiological and energetic investment strategies. In this framework optimal growth rate and the associated optimal physiology co-emerge from a set of interconnected constraints. We show that this optimization is intimately connected with how cells deal with degradation processes.

The focus on damage is likely to have far reaching implications for cellular physiology and ecology. For example, most cells are experiencing growth conditions that are far from optimal and cause them to grow at much slower rates, and cells only experience optimal growth conditions intermittently [20]. In very low energy environments, such as deep ocean sediments, the observed growth rate can be more than 10^6^ times slower than the maximum growth rate [6]. In these slow growth conditions metabolism is dominated by replacement processes. Thus, a very useful extension of our model here is to optimize growth under various values of the effective energy flux from the environment.

## V. AUTHOR CONTRIBUTIONS STATEMENT

CPK and DCK conceived of the study. KJ, CPK, SI-B, and DCK designed the research and developed the mathematical framework. KJ integrated and reformulated extant frameworks, spearheaded analytic calculations, and developed efficient calculation routines under the guidance of SI-B. RG checked calculations under the guidance of KJ and SI-B. KJ and CPK performed numerical analysis and developed the numerical code. KJ, SI-B, DCK and CPK produced the figures. All authors discussed the results and wrote the manuscript.

## VI. SUPPLEMENTARY INFORMATION

Here we provide a more detailed description of our theoretical framework and derivations. In summary, we use total available cellular energy and space as hard constraints under which we optimize the growth rate of a population by finding optimal allocations of metabolism and biosynthetic resources. In general we connect all of these features with cell size as a natural way to capture both changes across a division cycle and across diverse bacterial species.

### A. Energy Budgets and Growth

We begin with a perspective of how the total energy budget of an organism is used by the cell. The energetic effects on growth have been previously shown to be effectively modeled by the partitioning of total metabolic power, *B*, into biosynthetic and maintenance/repair purposes according to

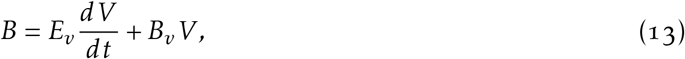

where *V* is cell volume size, *E_v_* is the energy to create a unit of volume (J V^-1^), and *B_v_* is the metabolic power to maintain an existing unit of cell volume (W V^-1^). It is typically the case that metabolic rate changes with organism size both within and across species according to a power law of the form

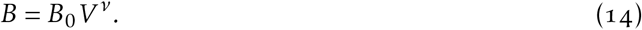

This budget forms the basic growth equation of our model because 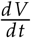 can be solved to give

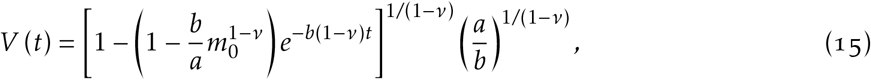

with *a_v_* = *B*_0_/*E_v_* (*g*^1-*v*^ s^-1^) and *b_v_* = *B_v_/E_v_*. Under the deterministic assumption that the cell divides exactly when the volume reaches *ϵ* times the initial volume, this equation can be used to find the time to double:

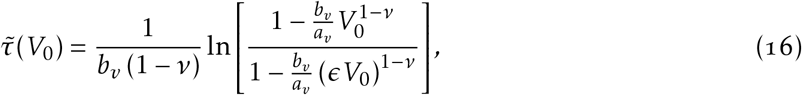

where *V*_0_ is the initial size of the cell, *ϵ* is 2 for symmetrically dividing cells, and where population growth rate is given by 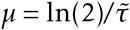.

### B. Macromolecule Scaling with Cell Size

#### 1. Macromolecule Scaling Functions

Beyond available energy, the abundances of macromolecules form the fundamental constraints of our model both in terms of available space, biosynthetic capacity, and the “replicative load” of a functional cell. Again we employ a size-based framework. Previous work [4, 5] using empirical data and theory has described the average number of ribosomes and proteins (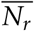 and 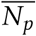 respectively, here the overhead bar denotes averaging over a cell cycle) as a function of *V_avg_*, the average volume of cell in a population, where these two macromolecules follow

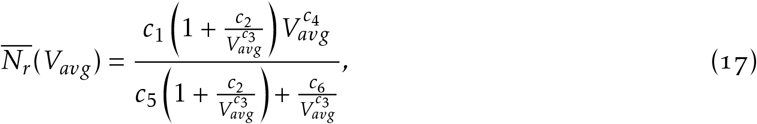

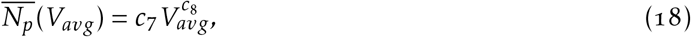

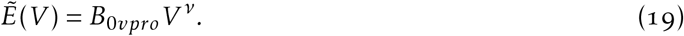

#### 2. Adapting to Particular Initial Size

For all of the optimizations in our model we need to understand what steady-state cell to expect for a particular size and set of energetic and physiological partitioning parameters. For adaptation to a particular size, we consider a cell starting from initial volume *V*_0_ and growing exponentially such that the volume is given by *V*(*t*) = *V*_0_*e^kt^*. In our model, we assume the cell divides exactly when volume doubles, thus division time *τ* = ln(2)/*k*. Thus, 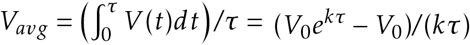. Replacing the value of *τ*,

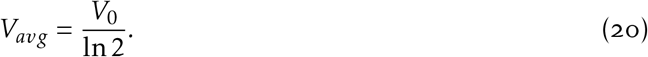

The biosynthesis partitioning (*γ*_0_) for an idealized cell in which ribosome pools exactly double in one division cycle can be calculated by setting *N_r_*(*c_r_,τ*) = 2*N_r_*(0) in Eq. 3a, thus,

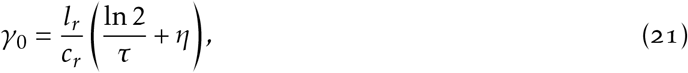

where *τ* can be obtained for a specific *V_avg_* through Eq.s 16 and 20. The initial number of ribosomes (*N*_*r*,0_) and proteins (*N*_*p*,0_) can be calculated from the average number of ribosomes and proteins by integrating Eq.s 3a and 3c over a division cycle. These turn out to be,

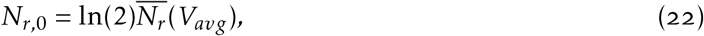

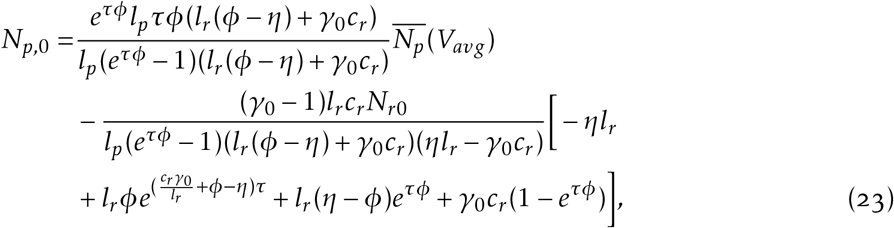

where, 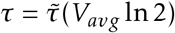. Also note that this partitioning ratio *γ*_0_ is simply used to calculate the initial ribosome and protein numbers for an idealized cell, and may not be the optimum partitioning ratio for cells of a given average size.

We assume that the instantaneous energy production in the cell depends more specifically on the number of functional proteins (which is proportional to the cell volume for the ideal case) rather than the cell volume itself. Thus, for the non-ideal case of variable accumulation of damaged pools, the instantaneous energy production will be given by,

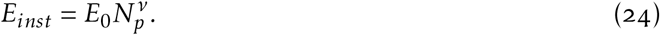

For cells of a given average volume, we can calculate *E*_0_ from the instantaneous energy production at birth, for an idealized cell with *N*_*p*,0_ proteins at birth. From Eq. 19,

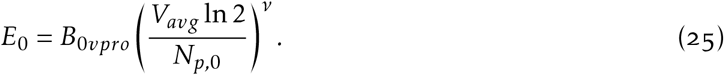

For simplicity, we assume that the total cell volume (*V*) is proportional to the sum of the volume occupied by functional and damaged pools of ribosomes and proteins, which we denote by *v.* The proportionality constant changes for species of different average sizes, and for a species of a given average volume, the value of *v* at birth is a constant given by,

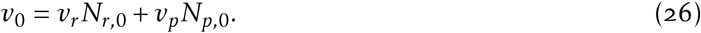

To get the above relation, due to a lack of data on size of damaged pools across species of different sizes, we have assumed that the idealized cells don’t have any accumulated damaged pools at birth. We have tested this assumption by comparing our model results for this case with the case in which the damaged pools at birth are exactly half the size of functional pools (thus, *v*_0_ is 1.5 times the value calculated through Eq. 26). We found that results for both cases are qualitatively identical, and only differ in absolute magnitude of values and the specific location of occurrence of transitions (Fig. 6, 7).

**FIG. 6:**
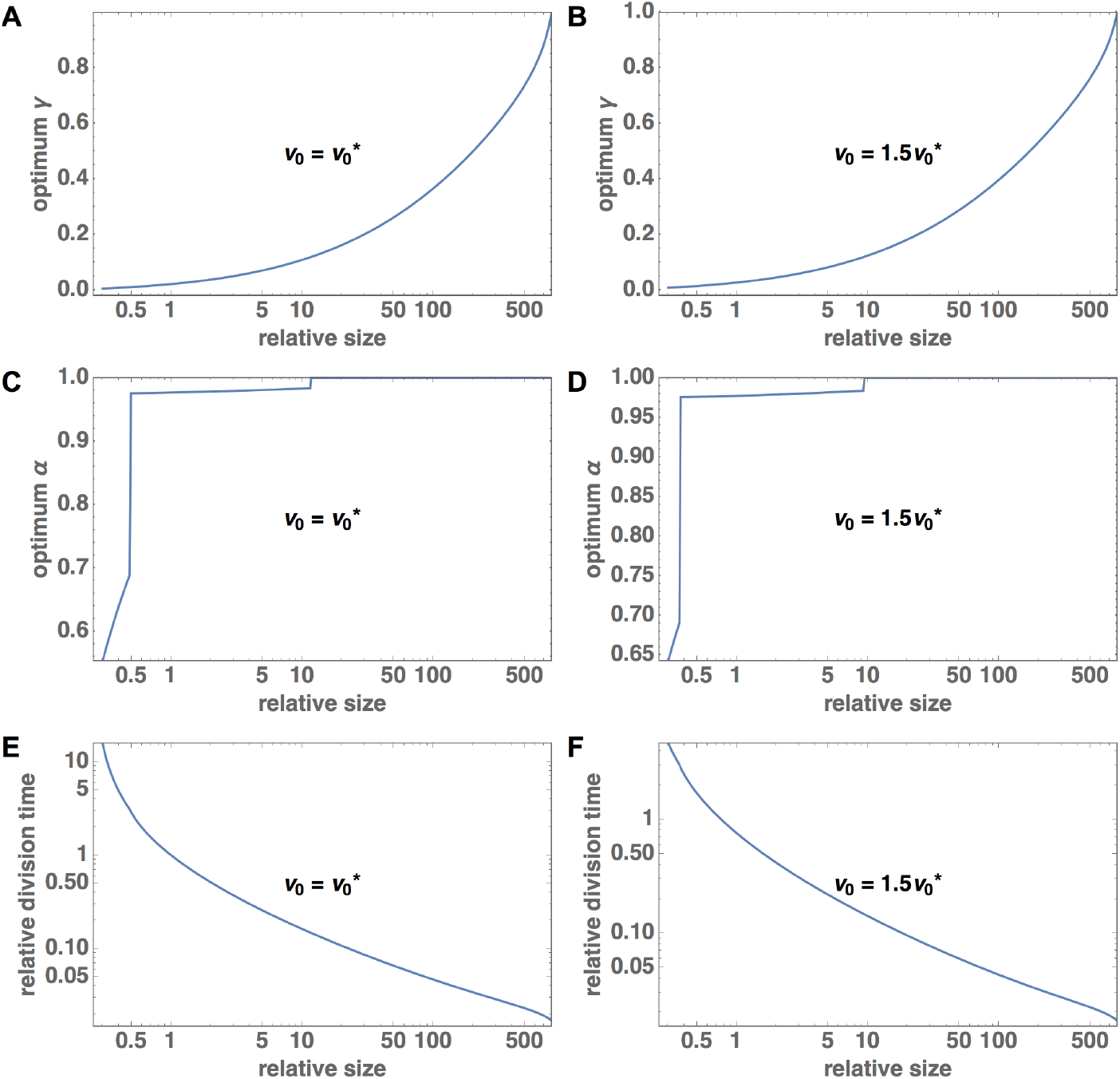
Scaling of optimum partitioning with cell size for different initial volume constraints. Variation of optimum A-B: *γ*, C-D: *α*, and E-F: division time with cell size. Degradation rate is kept constant at base degradation rate of *E. coli.* In A-C, degradation rate is normalized by the base degradation rate of *E. coli*, and in D-F, size is normalized by the size of *E. coli.* The plots on the left correspond to the case in which *v*_0_ is given by Eq. 26 and are identical to the corresponding plots in Fig. 4 (replicated here for comparison), while the plots on the right correspond to the case in which *v*_0_ is 1.5 times the value given by Eq. 26.

**FIG. 7:**
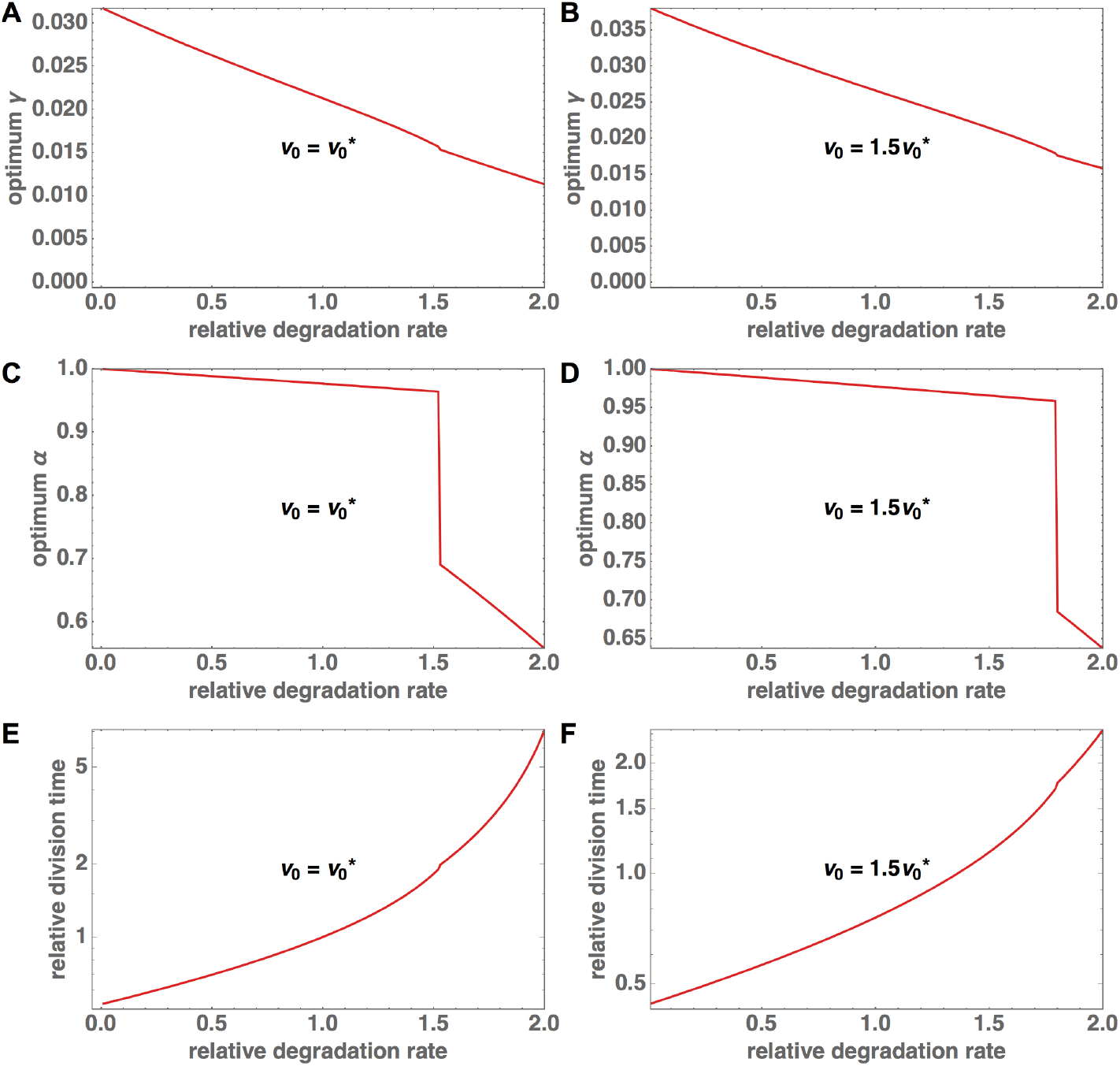
Scaling of optimum partitioning with degradation rate for different initial volume constraints. Variation of optimum A-B: *γ*, C-D: *α*, and E-F: division time with degradation rate. The plots on the left correspond to the case in which *v*_0_ is given by Eq. 26 and are identical to the corresponding plots in Fig. 3 (replicated here for comparison), while the plots on the right correspond to the case in which *v*_0_ is 1.5 times the value given by Eq. 26.

Of all the quantities derived in this section as a function of a given average cell size, only *v*_0_ and *E*_0_ enter our model as constants (for a given *V_avg_*), while the rest are simply used to derive these two quantities. These quantities are obtained from the average trends observed from multiple species, and may not accurately represent any particular species. Our model can be applied more accurately to a particular species by simply replacing the calculated values of *v*_0_ and *E*_0_ derived here by the known values for that species.

### C. Energy optimization

The total energy available to the cell during a cell cycle (*E_tot_*) is partitioned into growth and clearing damaged pools in the ratio *α* : 1 – *α*. In this section we discuss the strategy for using the thus allocated energy that minimizes division time for a given partitioning ratio *α*.

#### 1. Growth

The total ribosomes and proteins produced during a cell cycle are limited by the total energy available for growth during that cell cycle. We model this limitation by altering the polymerization rate *c_r_* (which is the primary factor in production of ribosomes and proteins), such that there is sufficient energy to sustain the production of ribosomes and proteins at this rate. To optimize growth, *c_r_* is adjusted such that it is the greatest possible rate for the given energy allocation. For a given division time *τ*, this is the maximum value (not greater than the physiological upper limit *c_r,max_*) that satisfies the inequality, *αE_tot_*(*c_r_,τ*) ≥ *E_req_*(*c_r_,τ*), where *E_req_* is the energy required for production of ribosomes and proteins, and is given by Eq. 7.

For the thus calculated value of *c_r_*, if *E_tot_* is strictly greater than *E_req_*, there will be excess energy left over from the portion allocated to growth. We allow this excess energy to go to waste and don’t transfer it to repair because our aim is to optimize the energy partitioning and transferring it to repair would make it indistinguishable from other choices of energy partitioning which result in the same effective energy distribution.

#### 2. Clearing Damaged Pools

The fraction 1–*α* of energy is spent by the cell in clearing out the volume occupied by damaged proteins and ribosomes. The cell spends *E_r,rem_* energy to remove one ribosome, which frees up volume *v_r_*. Thus, the volume freed per unit energy spent on damaged ribosomes is *v_r_*/*E_r,rem_*, and for damaged proteins is *v_p_*/*E_p,rem_*.

From empirical data, we find that *v_r_*/*E_r,rem_* > *v_p_*/*E_p,rem_*. Thus, cells can free up more volume with the same energy spent by prioritizing clearing damaged ribosomes. Thus, in our model, cells first spend all repair energy towards clearing damaged ribosomes. If all damaged ribosomes are cleared with allocated energy left over, this energy is now spent on clearing damaged proteins. If all damaged proteins are cleared too, we allow the remaining energy to go to waste instead of transferring it to growth because that would make it indistinguishable from other choices of energy partitioning which result in the same effective energy distribution.

Let

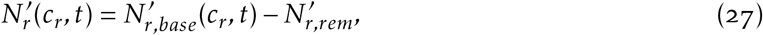

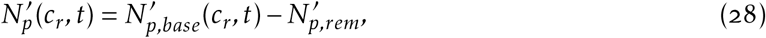

where 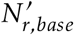 and 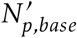 are the values of damaged ribosomes and proteins in the absence of any repair. Their expressions can be obtained from Eq.s 3b and 3d respectively. Then, according to the described strategy,

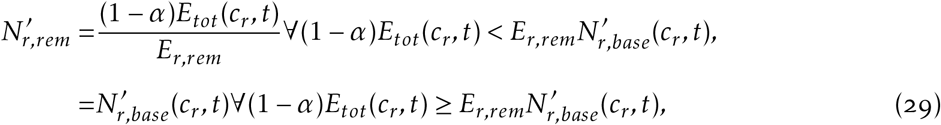

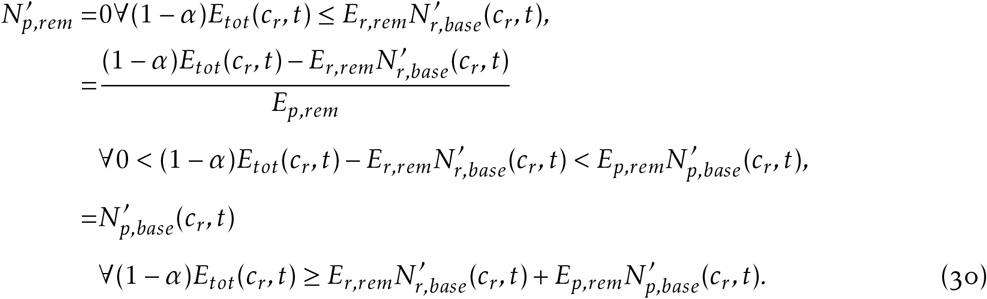

### D. Finding Division Time

In our model, the cells are constrained to divide exactly when their volume doubles. Since total volume is represented by the sum of volume occupied by ribosomes and proteins (both functional and damaged), thus division time is determined by when the sum of these quantities doubles. Also, these quantities are dependent on the polymerization rate, which in turn is determined by the energy allocated to growth which implicitly depends on both polymerization rate and division time. Thus, for a cell not in steady state, for a given division cycle that starts from a given number of functional and damaged ribosomes and proteins at birth, we find the polymerization rate and division time for that division cycle by solving the following two coupled non-linear equations,

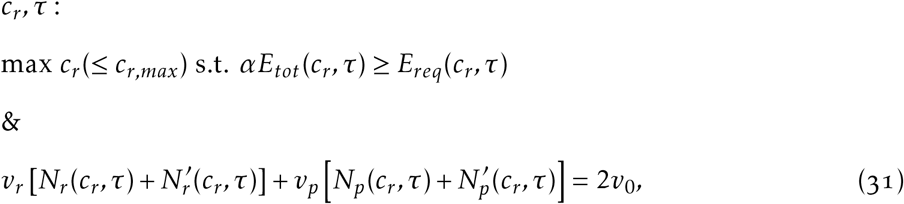

where the energy allocated to repair implicitly factors into 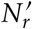 and 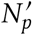 through Eq.s 29 and 30.

For cells starting from an arbitrary initial condition (an arbitrary composition of functional and damaged ribosomes and proteins), we need to use numerical methods to find the solution of these coupled equations on a case by case basis for each consecutive division cycle. For symmetrically dividing cells, the starting pool sizes for any division cycle are exactly half of the pool sizes at division from the previous division cycle. For cells in steady state, these equations can be reduced to a single equation with one variable which we have derived in the following section.

### E. Steady State

For a cell starting from an arbitrary initial condition, the polymerization rate and division time are determined by Eq. 31. These in turn determine the numbers of functional and damaged ribosomes and proteins contained in the cell at division. For simplicity, we assume division is perfectly symmetric with each daughter obtaining exactly half of each pool of functional and damaged ribosomes and proteins. Cells are said to reach steady state when the starting numbers of functional and damaged ribosomes and proteins in the daughter cells are the same as the starting numbers in the mother cell. Thus, these numbers exactly double during one division cycle in steady state.

In Eq. 3a, setting *N_r_*(*c_r_, τ*) = 2*N_r_*(0) gives us,

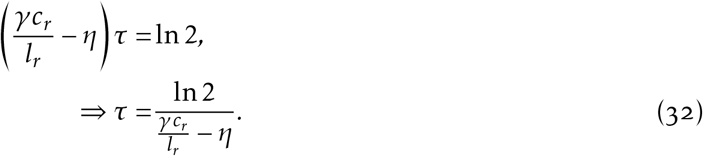

In Eq. 3c, setting *N_p_*(*c_r_,τ*) = 2*N_p_*(0) gives us,

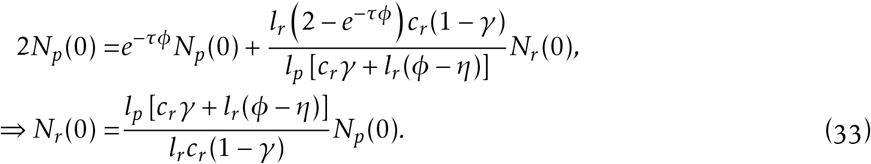

To find the total energy using Eq. 5, we need the instantaneous value of *N_p_* throughout the division cycle. Substituting the value of *N_r_*(0) from the above equation into Eq. 3c, we get,

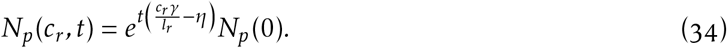

Now, substituting this into Eq. 5, we get,

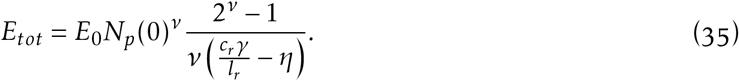

As previously defined, the energy required for growth *E_req_* is given by,

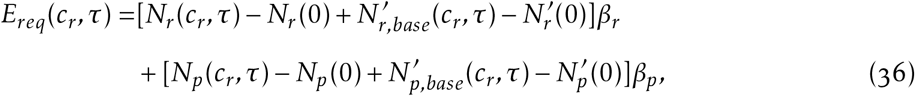

where, from Eq.s 3b and 27,

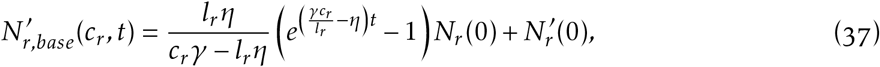

and from Eq.s 3d and 28, (38)

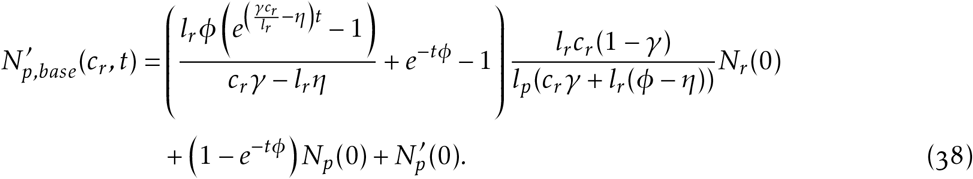

Factoring in the steady state simplifications (Eq.s 32 and 33), we get,

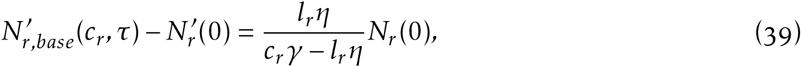

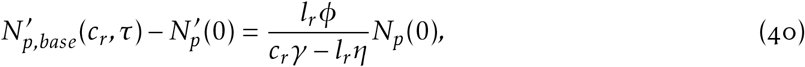

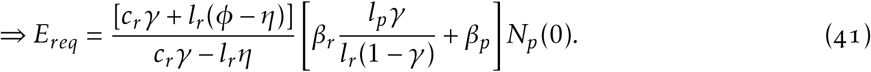

From the constraint *E_req_* ≤ *αE_tot_* and Eq. 35, we get the following constraint on *c_r_*,

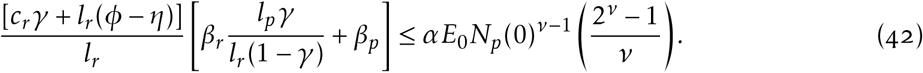

Now consider the following limiting value of *N_p_*(0), obtained by replacing *c_r_* by *c_r,max_* in the above constraint,

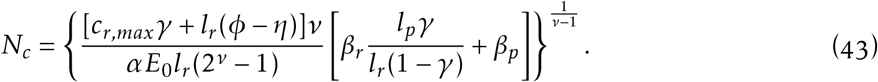

Since we are looking for the maximum value of *c_r_* (because that minimizes *τ* through Eq. 32) that satisfies the constraint Eq. 42, we have,

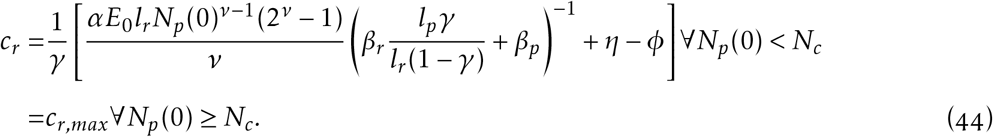

We can derive the expression for 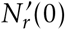 from Eq.s 27, 29, and 39, and using 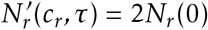. Thus, after replacing *N_r_*(0) through Eq. 33, we get,

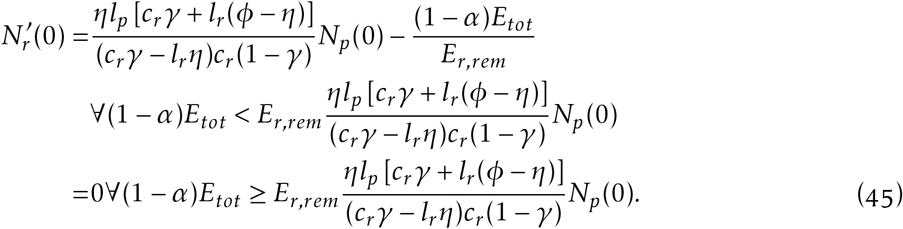

Similarly, from Eq.s 28, 30, and 40, and using 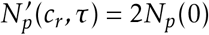, we get,

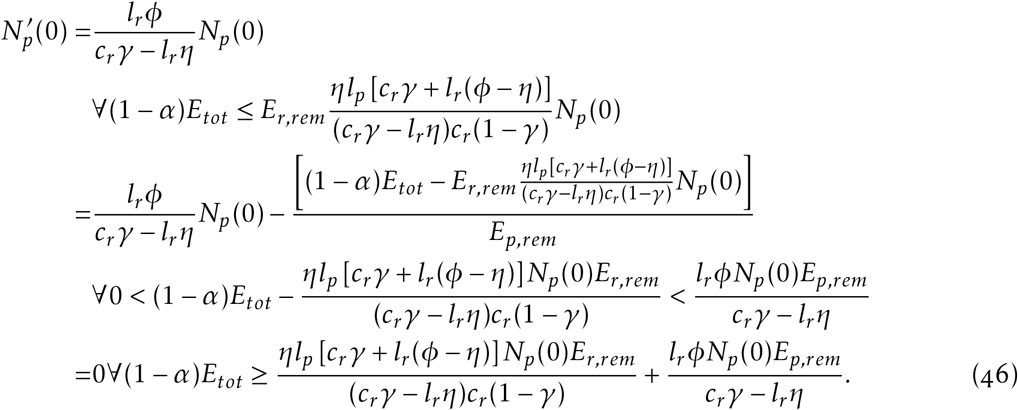

In our calculations for the steady state quantities, we have already incorporated the constraint that volume exactly doubles in a division cycle. To get the steady state quantities for a given initial size (or *v*_0_), we simply solve the following equation,

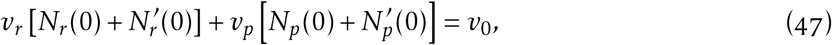

where, all quantities on the LHS can be expressed solely terms of *N_p_*(0) through Eq.s 33, 35, 44, 45, and 46. Numerically solving this gives us the steady state value of *N_p_*(0), which we can use to find the steady state division time through Eq.s 32 and 44.

### F. Limits on Partitioning Ratios

#### 1. Lower bound on γ (Fig. 2A i)

There exists a lower limit on fraction of biosynthetic capacity allocated to ribosome growth (*γ*), below which degradation would dominate over growth, leading to an exponential decay of ribosome numbers. In Eq. 3a, ensuring the exponent is positive for at least one *c_r_* value results in the following constraint,

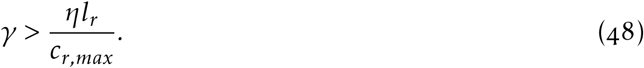

#### 2. Lower bound on a (Fig. 2A ii)

From the same constraint (obtained by ensuring exponential growth in Eq. 3a), for a given value of *γ*, the following gives the constraint on *c_r_*,

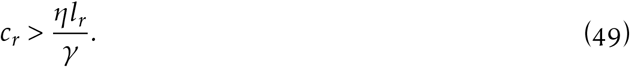

There exists a certain minimum energy requirement to sustain this level of polymerization rate. We can get a lower bound on *N_p_*(0) needed to generate this energy by replacing *c_r_* by Eq. 44 in the above inequality,

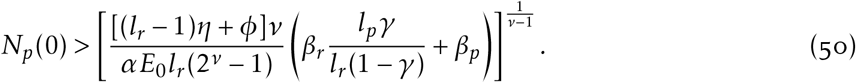

From here, the lower bound of *N_r_*(0) can be found using Eq. 33. The lower bounds of 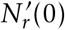 and 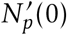 are o as long as sufficient energy is available. Since the total volume of a cell is limited, the total volume occupied by the lower bounds of ribosome and protein numbers must not exceed this limit. From Eq. 33, 47, and 50, we have,

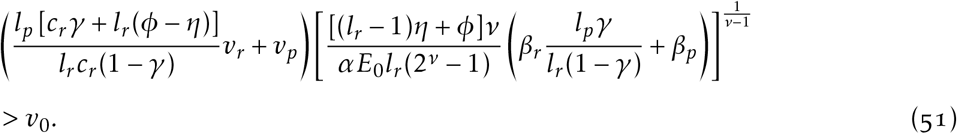

Now consider the term,

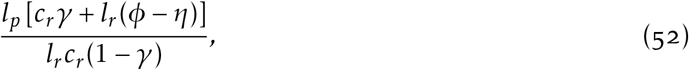

which is a strictly monotonically decreasing function of *c_r_*. Thus, its maximum value is attained at the minimum value of *c_r_*. Thus, we can preserve the above inequality by replacing *c_r_* with its minimum value from Eq. 49, giving us,

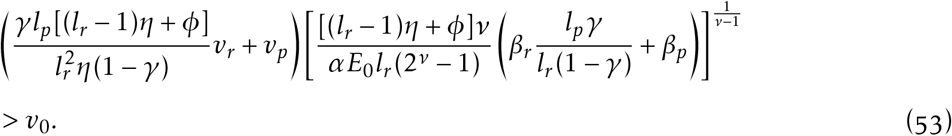

Thus, for a given value of *γ*, the lower bound on *α* is,

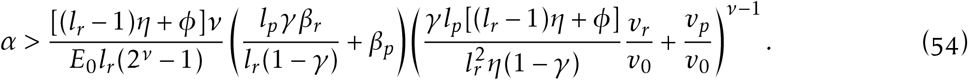

#### 3. Maximum γ beyond which α = 1 is not viable (Fig. 2A iii)

For values of *γ* above a certain threshold, allocating all energy to growth (*α* = 1) is no longer viable. If no energy is allocated to repair, the damaged pools keep growing every subsequent cell cycle, ultimately making growth non-viable.

Here, we find the analytical solution for this threshold for the case in which *η* = *ϕ* (which is true for all results presented in the main text), but the solution for the general case can be derived analogously. For the case when no energy is allocated to removal of damaged pools (*α* = 1), after writing all terms in Eq. 12 in terms of *N_p_*(0) (see section VI E), we get,

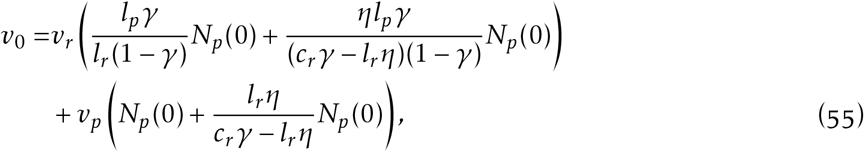

where, from Eq. 11,

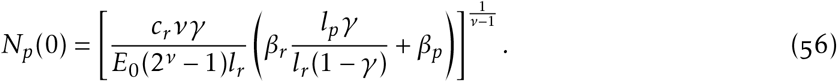

Thus, the RHS of Eq. 55 is a function of *c_r_* given by,

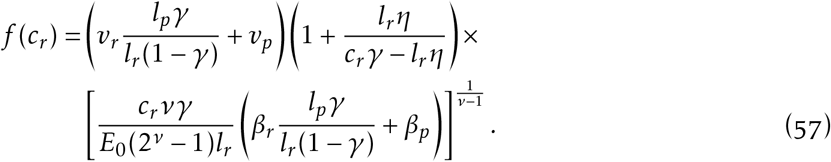

We are looking for a solution for *c_r_* between *l_r_η/γ* (Eq. 49) and *c_r,max_* that satisfies *f* (*c_r_*) = *v*_0_. As *f*(*c_r_* → *l_r_η/γ*) → ∞, if the minimum of *f* is greater than *v*_0_, there would be no solution possible, while minimum being less than or equal to *v*_0_ would guarantee a solution. We find the local extrema by setting *f*′(*c_r_*) = 0, and find that the only local extremum in the given domain lies at *c_r_* = *νl_r_η/γ*. This must be the minimum because *f* is decreasing as *c_r_* increases from *l_r_η/γ*. Thus, the minimum value of *f* is,

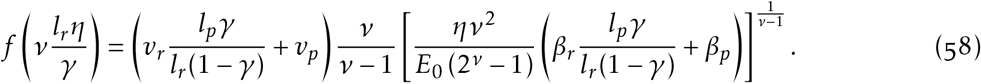

This is a strictly increasing function of *γ*, which tends to ∞ as *γ* → 1. Thus, there exists a threshold on *γ* above which no solution exists for *f*(*c_r_*) = *v*_0_. This threshold is given by solving the following implicit equation:

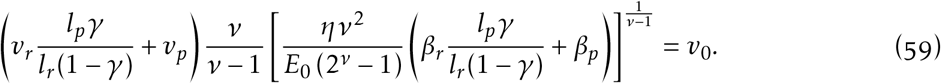

#### 4. First upper bound on a (Fig. 2A iv)

As *γ* increases beyond the above threshold, less biosynthesis is allocated to proteins reducing energy production, thus more energy fraction needs to be allocated to removal of damaged ribosomes in order to maintain viable growth (i.e., max value of *α* decreases). Thus, there exists an upper bound on the value of *α* above which growth is not viable. This upper bound can be found using a similar procedure to that used in the last subsection. For this case, Eq. 12 becomes (obtained by simply adding the removal of damaged ribosomes in Eq. 45 to Eq. 55, and using Eq. 35 for the expression for *E_tot_*),

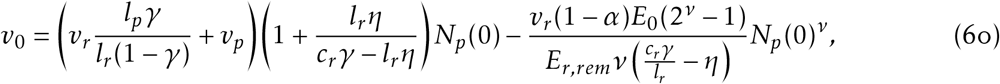

where, from Eq. 11,

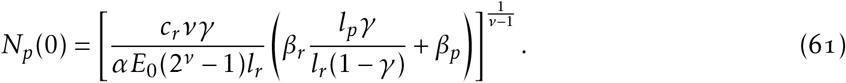

Thus, we need to solve for *f*(*c_r_*) = *v*_0_, where *f* for this case is given by,

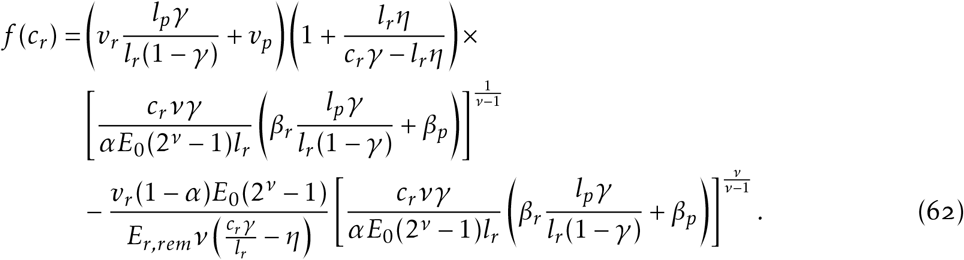

Rearranging the terms, we get,

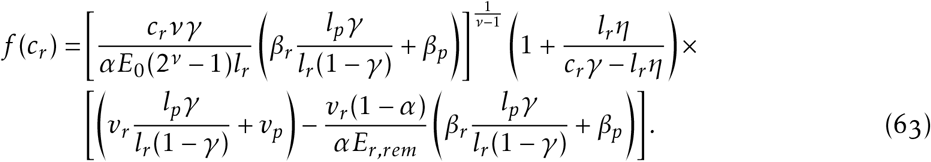

This new *f* is just a constant times the *f* from the previous case, thus the location of minimum is the same, at *c_r_* = *νl_r_η/γ*. Similar to the previous case, we get the upper bound on *α* for a given value of *γ* by solving the implicit equation *f*(*νl_r_η/γ*) = *v*_0_ (a solution for steady state is guaranteed as long as *f*(*νl_r_η/γ*) ≤ *v*_0_). Thus, we get the following implicit equation for the maximum value of *α* for viable growth for a given *γ*,

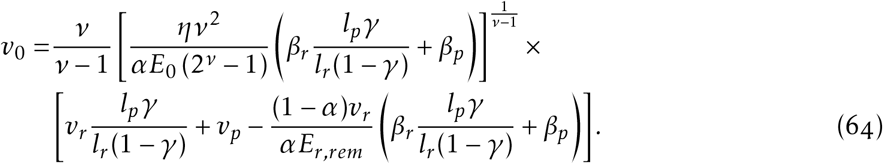

#### 5. First transition in upper bound on a (Fig. 2A v)

As *γ* increases, at a certain value of *γ*, the maximum *α* is such that all damaged ribosomes are repaired. On increasing *γ* further, the additional energy needs to be spent on repairing damaged proteins instead. At the *γ* value corresponding to this transition, we will have 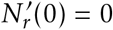 for the upper bound on *α* given by 64. Thus, from Eq. 45,

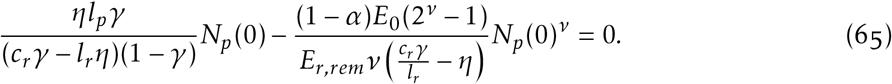

Now, replacing the value of *N_p_*(0) from Eq. 61, and replacing *c_r_* by *νl_r_η/γ* from the previous subsection, we get the following result for the value of *α* at this transition point,

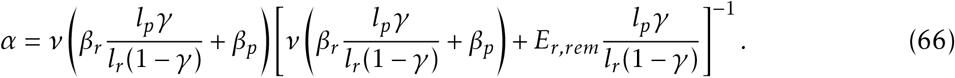

Replacing this in Eq. 64, we get the following implicit equation for the *γ* value at which this transition between first upper bound (all energy spent on ribosome repair) and second upper bound bound (damaged ribosomes fully repaired, remaining energy spent on protein repair) occurs,

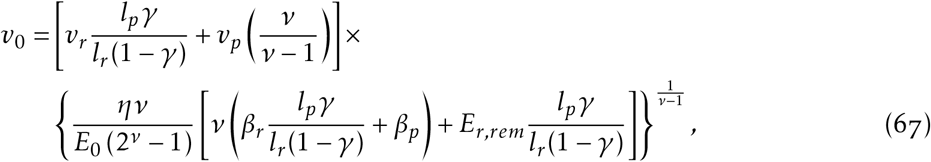

and, at this value of *γ*, the value of *α* is given by Eq. 66.

#### 6. Second upper bound on a (Fig. 2A vi)

As *γ* is increased beyond the above transition point, the minimum amount of repair required for viable growth is so high that all damaged ribosomes need to be fully repaired, and remaining repair needs to go to damaged proteins. In this regime, after writing all terms in Eq. 12 in terms of *N_p_*(0) (see section VI E), we get,

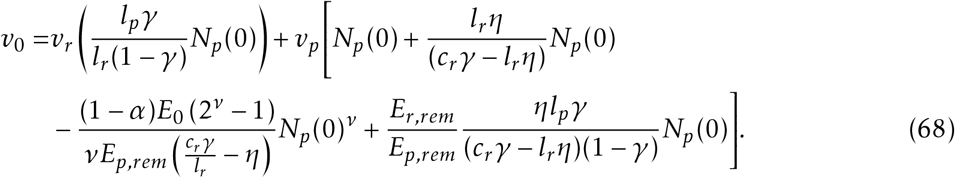

Replacing the value of *N_p_*(0) by Eq. 61, and changing variables from *c_r_* to *x* = *c_r_γ*/(*l_r_η*), the above equation can be expressed as *v*_0_ = *f* (*x*), where *f* (*x*) is,

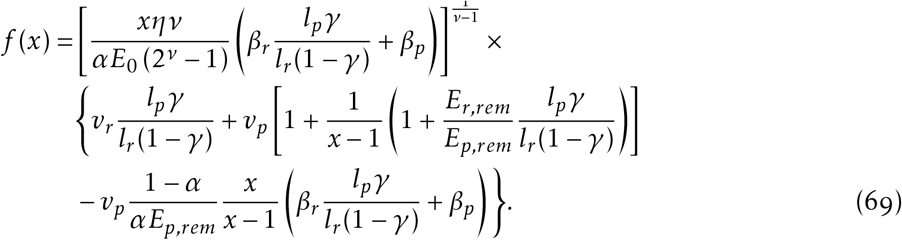

Proceeding similar to the previous subsections, we find the minimum of *f*(*x*) by setting its derivative w.r.t. *x* to 0. After some simplification, we get,

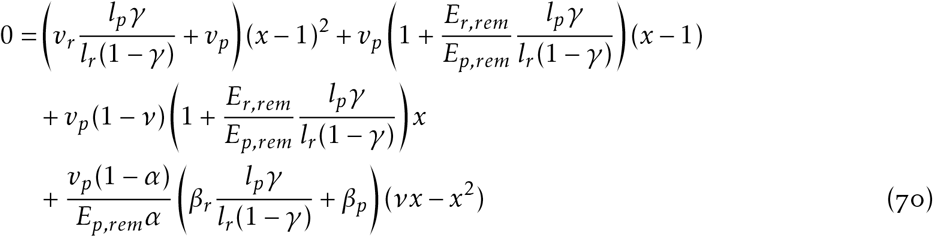

Labeling the RHS as *g*(*x*), we need to find the root of *g*(*x*) that lies above 1 due to constraint Eq. 49. Consider,

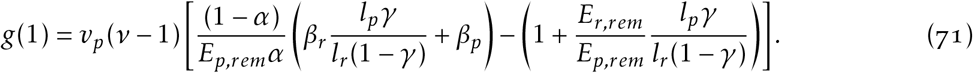

For the given values of *E_r,rem_, E_p,rem_*, *β_r_*, *β_p_*, and *v, g*(1) < 0 irrespective of the value of *γ* as long as *α* > 0.5. In the next subsections, we have calculated the *α* value at which all damaged pools are fully cleared, and this value happens to lie above 0.5 for this range of *γ* values. Since the *α* values in consideration in this subsection must lie above this (because all damaged proteins are not fully cleared), so in this subsection we are not concerned about what happens for *α* ≤ 0.5. Also, the coefficient of *x*^2^ in *g*(*x*) is,

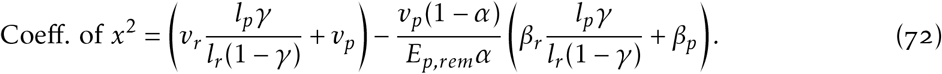

For the given values of constants, this is always positive for *α* > 0.5, irrespective of the value of *γ*. Thus, *g*(*x* → ±∞) → ∞, and *g*(1) < 0. Thus, the smaller root of the quadratic polynomial *g*(*x*) lies below 1, while the larger root lies above 1. Thus, the point of minimum we are searching for is at the larger root, which for a given value of *α* and *γ*, is given by,

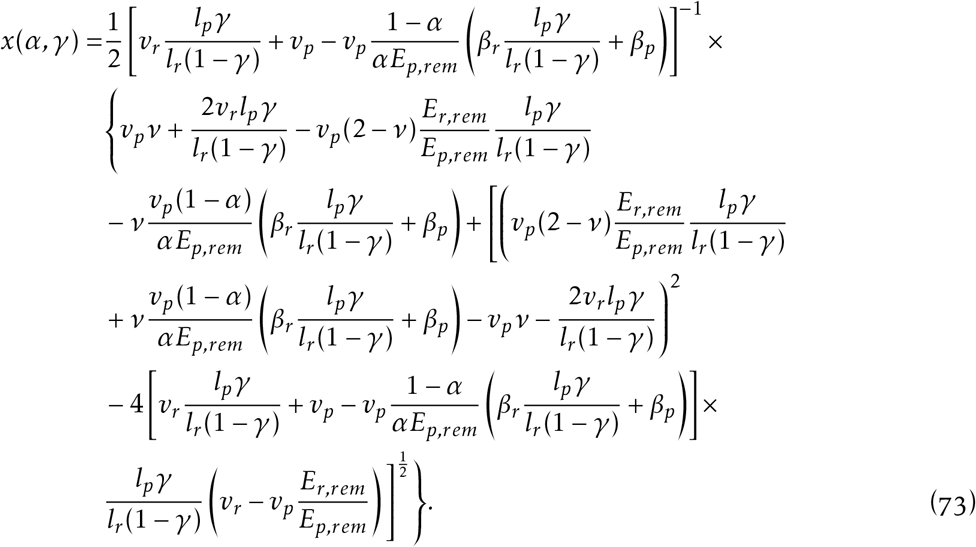

Replacing this back into the expression for *f*(*x*), for a given *γ* in this range, we can find the maximum *α* for viable growth through the following constraint:

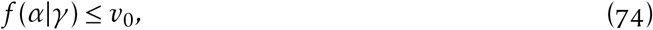

where,

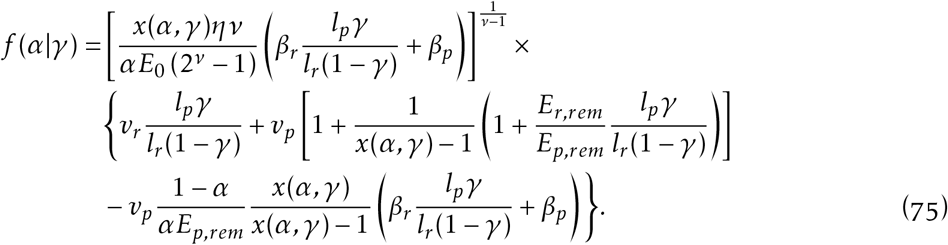

For the range of *γ* values under consideration, *v*_0_ = *f*(*α*|*γ*) has two roots, *α*_1_ < *α*_2_, such that *v*_0_ ≥ *f*(*α*|*γ*) for all *α* ≤ *α*_1_ or *α* ≥ *α*_2_.

Thus, for a given value of *γ*, growth is viable for *α_min_* < *α* ≤ *α*_1_ (where *α_min_* is the lower bound, given by Eq. 54), not viable for *α*_1_ ≥ *α* ≥ *α*_2_, then viable again for *α*_2_ ≤ *α* ≤ *α*_3_, where *α*_3_ represents the minimum energy allocated to removal of damaged pools to ensure all damaged ribosomes are removed.

When *α* ≥ *α*_2_, the energy allocated to growth is enough to sustain large number of proteins, which in turn results in large energy availability, enough to ensure growth remains viable. But, when *α*_1_ ≥ *α* ≥ *α*_2_, there in not enough energy allocated to either growth or removal of damaged pools, making growth not viable. For *α* ≤ *α*_1_, the energy allocated to removal of damaged pools becomes large enough to ensure enough damaged pools are removed to sustain growth.

To calculate *α*_3_, we have 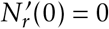, thus (from Eq. 45 and 35),

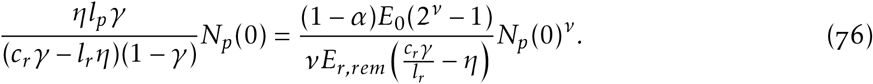

On replacing the value of *N_p_*(0) from Eq. 61 and simplifying, we get

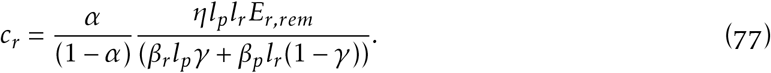

Also, from Eq. 12,

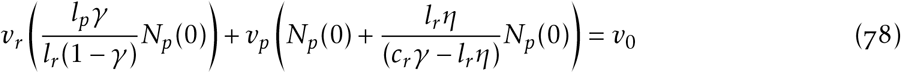

Plugging in the calculated value of *c_r_*, and *N_p_*(0) from Eq. 61, and simplifying, we get the following implicit equation for *α*_3_ (blue curve) for a given *γ*:

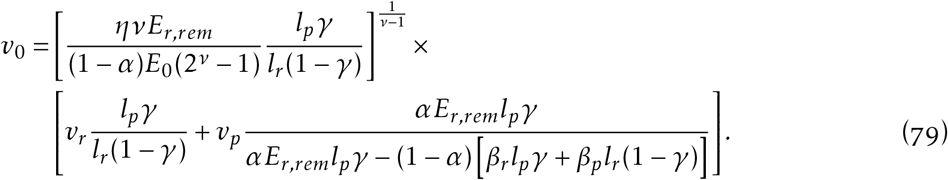

As *γ* increases, *α*_2_ eventually becomes greater than *α*_3_. Beyond this point, there is no solution for *α* > *α*_1_, thus *α*_1_ becomes the maximum *α* for viable growth.

#### 7. Second transition in upper bound on a (Fig. 2A vii)

Beyond a certain value of *γ*, the minimum energy fraction required to be allocated to clearing damaged pools is sufficient to completely remove all damaged ribosomes and proteins. For a given *γ*, if we set the value of 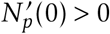, we get the lower bound on *α* for damaged proteins to not be fully removed. From Eq. 46 and 35, we have,

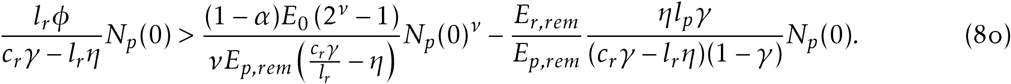

Rearranging the terms, we have,

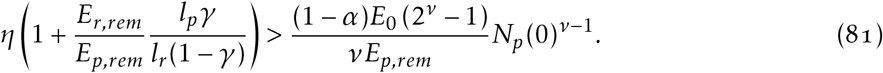

Replacing *N_p_*(0) by Eq. 61,

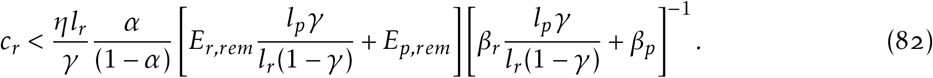

Now, converting from *c_r_* to *x* = *c_r_γ*/(*l_r_η*),

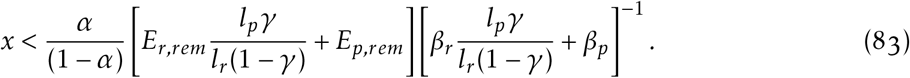

When the location of minimum for *f*(*x*) from previous subsection lies below this limit, the upper bound given by the previous subsection is viable. But, when that minimum crosses this limit, that minimum goes out of bounds for the case in which all damaged proteins are not fully removed, and the new minimum occurs at this limit. At this limit, *α* is exactly sufficient to fully remove all damaged pools without wasting any energy. For a given value of *γ*, this transition occurs at the value of *α* obtained by equating *x* in Eq. 73 with the RHS of Eq. 83. Thus, for a given value of *γ*, we have the following implicit equation for *α*,

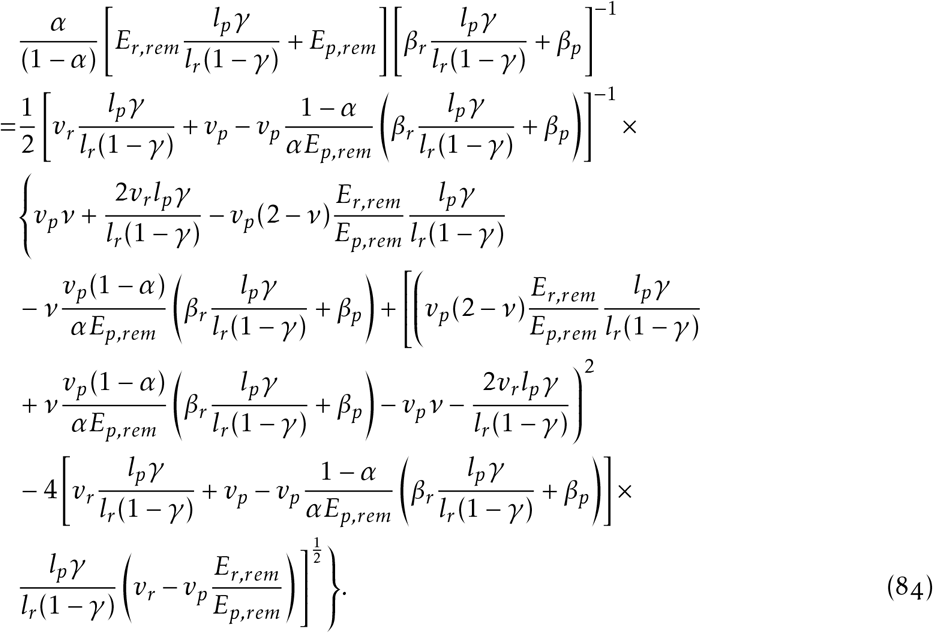

By solving this equation for *γ* in the range of values for the bound calculated in previous subsection, we found that this transition *α* is consistently greater than 0.5 throughout the range, thus validating our claim in the previous subsection that *α* ≤ 0.5 case doesn’t need to be considered.

Now. for the case in which all damaged pools are fully cleared. Eq. 12 becomes (from Eq 33).

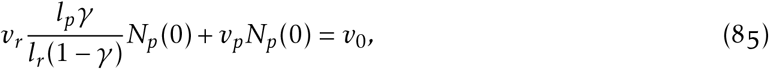

thus,

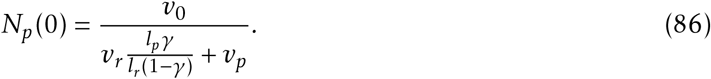

Eq. 81 reverses inequality for the case in which all damaged pools are fully cleared. Equating LHS with RHS gives the maximum value of *α* for which a solution for all damaged pools cleared exists. Replacing the value of *N_p_*(0) with the above equation, we get,

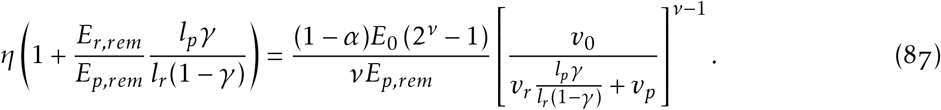

Thus, the maximum value of *α* for a given *γ* is given by,

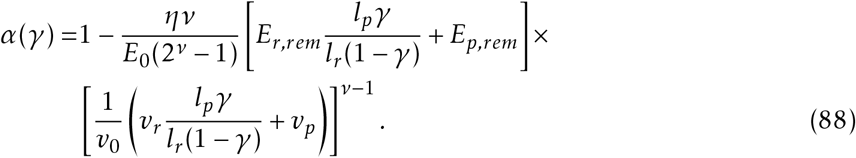

Thus, the value of *γ* at this transition point is given by the following implicit equation,

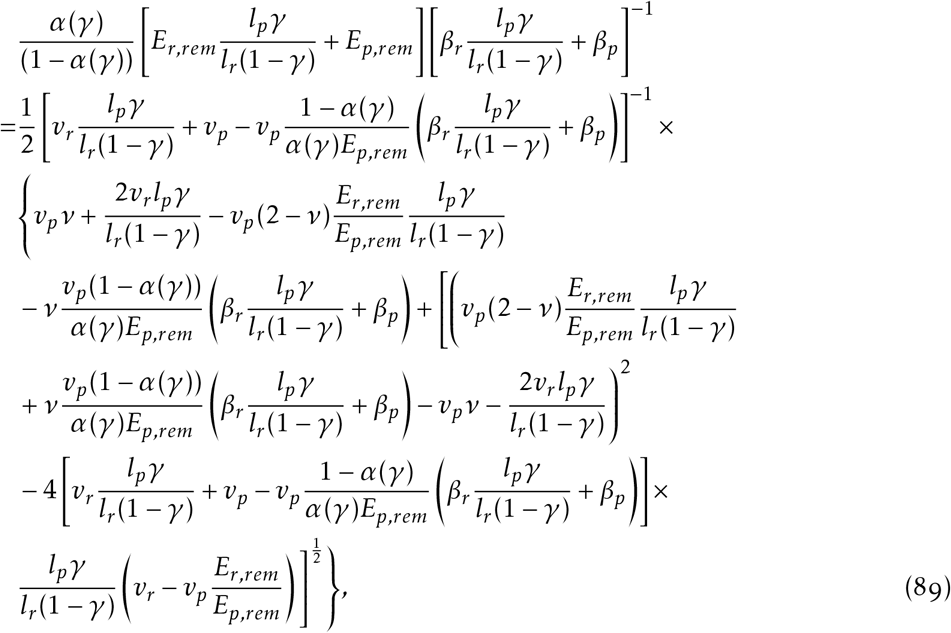

where *α*(*γ*) is given by Eq. 88. For the thus solved value of *γ*, the value of *α* at the transition point can also be found through Eq. 88.

#### 8. Third upper bound on a (Fig. 2A viii)

For values of *γ* beyond the above threshold, all damaged pools must be cleared to sustain growth. As derived in the previous subsection, for a given value of *γ*, the maximum value of *α* that allows this is given by Eq. 88.

#### 9. Upper bound on γ (Fig. 2A ix)

Growth is no longer possible beyond the *γ* value at which the upper and lower bounds on *α* intersect. This *γ* value can be found by equating Eq. 54 (after simplifying using *η* = *ϕ*) with Eq. 88. Thus, the maximum viable *γ* value is given by the implicit equation,

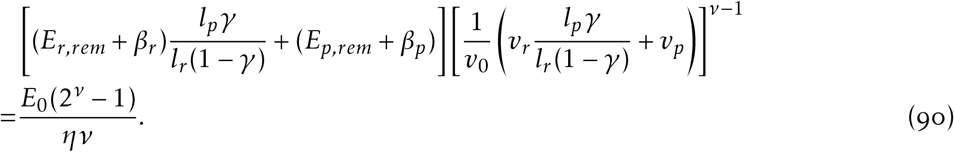

